# Harnessing Agent-Based Modeling in CellAgentChat to Unravel Cell-Cell Interactions from Single-Cell Data

**DOI:** 10.1101/2023.08.23.554489

**Authors:** Vishvak Raghavan, Yue Li, Jun Ding

**Author notes:** Corresponding authors, YL; JD.

## Abstract

Understanding cell-cell interactions (CCIs) is essential yet challenging due to the inherent intricacy and diversity of cellular dynamics. Existing approaches often analyze global patterns of CCIs using statistical frameworks, missing the nuances of individual cell behavior due to their focus on aggregate data. This makes them insensitive in complex environments where the detailed dynamics of cell interactions matter. We introduce CellAgentChat, an agent-based model (ABM) designed to decipher CCIs from single-cell RNA sequencing and spatial transcriptomics data. This approach models biological systems as collections of autonomous agents governed by biologically inspired principles and rules. Validated against seven diverse single-cell datasets, CellAgentChat demonstrates its effectiveness in detecting intricate signaling events across different cell populations. Moreover, CellAgentChat offers the ability to generate animated visualizations of single-cell interactions and provides flexibility in modifying agent behavior rules, facilitating thorough exploration of both close and distant cellular communications. Furthermore, CellAgentChat leverages ABM features to enable intuitive in silico perturbations via agent rule modifications, pioneering new avenues for innovative intervention strategies. This ABM method empowers an in-depth understanding of cellular signaling interactions across various biological contexts, thereby enhancing in-silico studies for cellular communication-based therapies.

## INTRODUCTION

Cells, as the fundamental units of most life forms, are dynamic and continuously adapt their states in response to changes in their microenvironment^1,2^. In multicellular organisms, cellular activities rely on the coordination of cell-cell interactions (CCIs) across different cell types and tissues^3–6^. CCIs are critical in numerous biological processes, including cell differentiation^7^, disease pathogenesis^8^, and immune responses^9^. Signaling molecules, such as ligands and receptors, facilitate cellular interactions and influence cell behavior, leading to dynamic changes in cellular functions^3^. By studying these interactions, we can gain insights into the mechanisms underlying biological processes, such as organ development and tumor growth, ultimately informing the study of various disorders and diseases and aiding in the identification of potential diagnostic and therapeutic strategies^9^.

However, inferring CCIs in biological processes is challenging due to cellular heterogeneity in most biological systems^10^. The advent of single-cell sequencing, particularly single-cell RNA-sequencing (scRNA-seq) and spatial transcriptomics measurements like 10x genomics Visium^11^, Slide-Seq^12^, MERFISH^13^, SeqFISH+^14^, Stereo-seq^15^ and Xenium^16^ offers unparalleled opportunities to investigate CCIs in diverse biological processes. Numerous tools have been developed in recent years for CCI inference by integrating known ligand-receptor (LR) pairs with gene expression data from scRNA-seq data, including state-of-the-art methods such as CellPhoneDB^17^ and CellChat^18^. Despite advancements, earlier methods and versions primarily rely on unimodal scRNA-seq data, which restricts their capacity to infer CCIs due to the absence of spatial distance information between cells^17–19^. As CCIs in tissue environments are heavily influenced by spatial structures and can occur at both long and short ranges^20^, integrating spatial information is essential for studying endocrine and paracrine signaling, respectively. Thus, several recent methods, such as COMMOT^21^, NICHES^22^ and CellChat v2^23^, have been developed for inferring cellular interactions from single-cell spatial measurements.

These existing methods still face significant drawbacks regardless of whether they integrate spatial information or not. Specifically, existing methods such as CellPhoneDB and CellChat depend on mean expression values computed from single-cell clusters and therefore focus on cell population levels, inferring CCIs between cell populations (clusters). Such approaches fail to fully exploit the single-cell resolution provided by the original measurements, resulting in a loss of the intricate signaling patterns that exists between cells. NICHES, COMMOT, and Scriabin^24^ are newer methods for analyzing LR interactions at the single-cell level. COMMOT applies collective optimal transport with spatial data to infer CCIs yet cannot use scRNA-seq data alone. Conversely, Scriabin assesses CCIs using solely scRNA-seq data, lacking spatial insights. Additionally, no method yet models both long-range and short-range interactions effectively. These limitations hamper the accurate discovery of CCIs in various biological processes^25^. Finally, CCIs play crucial roles in the pathogenesis of many diseases^8^, and the analysis of these interactions holds the potential for developing effective therapeutics. However, existing methods like COMMOT and scSeqComm^26^, despite their ability to capture the effects of CCIs on downstream genes, do not support in-silico perturbation to utilize these impacts for exploring potential treatments, such as receptor antagonists.

Agent-based model (ABM) simulates the actions and interactions of autonomous agents, through which it helps understand the behaviour and outcomes of a complex system^27^. In ABMs, agents are typically individuals or entities endowed with specific characteristics, behaviors, and rules. These agents can interact with each other and their environment, often leading to emergent behaviors and patterns at the macroscopic level. Therefore, ABM allows for a straightforward yet effective depiction of cellular interactions, embracing the concept of emergence. In the context of this study, this translates to intricate CCI patterns evolving from basic, local rules applied to individual cells’ interaction behaviors. This ABM methodology starkly diverges from existing top-down methods described above that infer interactions through global statistical correlations, often overlooking localized cellular dynamics and individual properties. Although ABMs hold great potential, to our knowledge, they have not been fully utilized for inferring CCIs from single-cell data. Due to their unique characteristics, ABMs are well-suited for exploring complex CCIs and their impact on spatial and temporal cellular dynamics in biological systems. First, ABMs capture emergent phenomena arising from collective agent interactions, revealing insights into cellular community function^27,28^. Second, they represent heterogeneous cells as agents with different statuses, enabling accurate modeling of diverse cell populations and capturing variations in cell types, states, and behaviors critical for understanding biological processes^27,28^. Third, ABMs, like many machine learning approaches, offer explicit integration of temporal and spatial information, a crucial capability for studying complex biological processes such as tissue development, immune responses, and tumor growth^27,28^. Lastly, ABMs facilitate the straightforward and effective adjustment of agent rules, enabling perturbations in agent behavior. This allows for predictions of alterations in cellular interactions and facilitates the search for potential therapeutics, like receptor antagonists^29^, based on modified cell agent behavior rules, such as receptor blocking.

In this study, we introduce CellAgentChat, a comprehensive ABM framework for inferring and visualizing CCIs from scRNA-seq and spatial transcriptomics data. Leveraging the inherent advantages of ABMs, CellAgentChat offers versatile functionalities for CCI inference and visualization. By adopting an agent-based perspective, CellAgentChat seamlessly integrates spatial and biological data, providing a more precise and comprehensive understanding of cellular interaction dynamics at both individual cell and system levels. Moreover, CellAgentChat enables in silico perturbations and in-depth analysis of the downstream effects of cellular interactions on gene expression and facilitates the identification of potential therapeutic receptor targets. Notably, the flexible framework of CellAgentChat allows for diverse functionalities by adjusting the behavior rules associated with cell agents. This feature enables the identification and modeling of long-range and short-range LR interactions. Finally, CellAgentChat has the capability to analyze temporal dynamics, shedding light on how CCIs influence cellular behavior over short time frames. Our evaluation of CellAgentChat, using several publicly available single-cell sequencing and spatial transcriptomics datasets, reveal superior CCI inference accuracy compared to other benchmarked methods. Together, we introduce a new paradigm of framework based on ABMs for future advancements in the study of cellular interactions implicated in complex biological processes.

## RESULTS

### Overview of CellAgentChat

CellAgentChat constitutes a comprehensive ABM framework integrating gene expression data and existing knowledge of signaling LR interactions to compute the probabilities of CCI. Figure 1 schematically outlines the CellAgentChat methodology. Utilizing the principles of ABM, we characterize each cell agent through various cellular attributes (states), including cell identities (e.g., cell type or clusters), gene expression profiles, LR universe and spatial coordinates (Fig. 1a).

**Figure 1:**
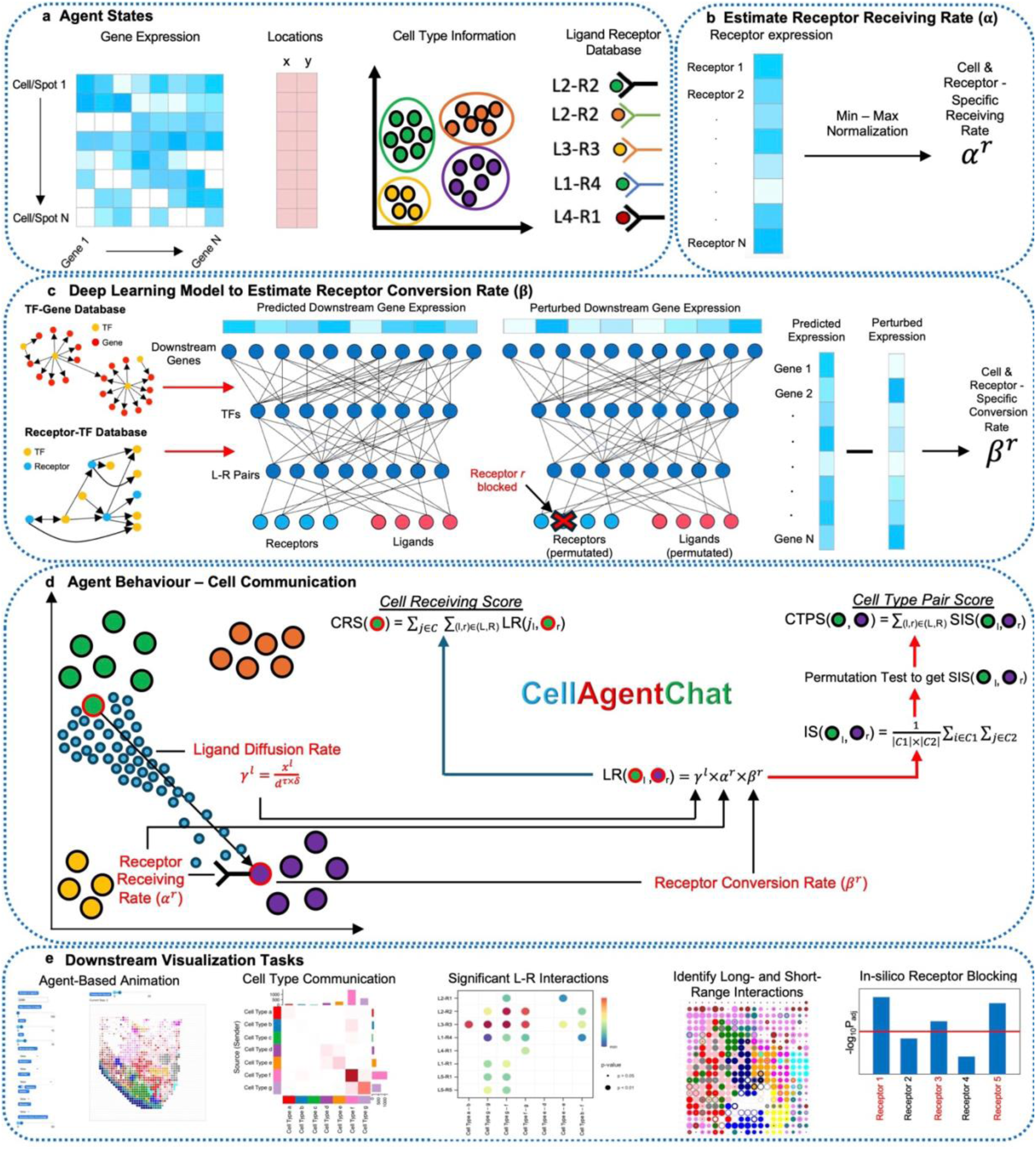
Overview of CellAgentChat. **a**, CellAgentChat requires the following agent states for each cell: gene expression derived from scRNA-seq data, its cell type/cluster information, and the ligand-receptor database (by default uses CellTalkDB). Optionally, users can provide spatial coordinates of each agent (from spatial transcriptomics data). **b,** The receptor receiving rate is the probability that an interaction is received by a receptor, calculated by non-parametric min-max normalization of the receptor expression. **c,** The inputs of our deep learning model are the ligand and receptor’s gene expression, and the outputs are the gene expression of all genes. The model incorporates prior knowledge of TF-receptor and TF-gene interactions from pre-existing databases to create a partially connected feedforward network. For a receptor, we compute the perturbed expression by permuting the feature variables of each receptor and scaling down the expression, thereby simulating receptor blocking. The receptor conversion *β*^*r*^of the receptor is estimated from the total change of the predicted expression after the perturbation. **d,** Calculating the ligand-receptor score between two cells for a specific ligand-receptor pair requires the ligand diffusion rate *λ*^*l*^, the receptor receiving rate *α*^*r*^, and the receptor conversion rate *β*^*r*^ . If spatial data is provided, the ligand diffusion rate *λ*^*l*^incorporates the Euclidean distance *d* between the two cells. The *τ* hyperparameter controls the degrees of spatial freedom for the distance (default *τ* =2). The *δ* hyperparameter (default *δ*=1) controls the decay rate of ligands and can be used to prioritize long-range (0 < *δ* < 1) or short-range (1 < *δ* < 10) interactions. *x*^*l*^ is the normalized ligand expression. Using the LR score, we can calculate the interaction score between two clusters/cell types and the cell receiving score (CRS) for a cell. After a permutation test, significant interactions are identified and used to calculate the cell type pair score (CTPS) between two clusters. **e,** CellAgentChat enables agent-based animation, visualizing cell type communication, finding significant LR pairs, identifying long-range and short-range interactions and in silico-permutations to simulate blocking receptors to identify the most perturbed downstream genes to discover therapeutic drugs.

We quantify cellular interactions between sender and receiver cells based on the number of ligands secreted by the sender cells and subsequently received by the receiver cells. This process hinges upon three interrelated components: ligand diffusion rate (*λ*^*l*^), receptor receiving rate (*α*^*r*^), and receptor conversion rate (*β*^*r*^) (Fig. 1b, c, d).

The initial step in our approach involves the estimation of the receptor receiving rate (*α*^*r*^), which signifies the fraction of secreted ligands received by the receptor on the receiver cell (Fig. 1b). We then leverage a neural network to evaluate the receptor conversion rate (*β*^*r*^), which measures the potency of a cellular interaction by studying its effect on downstream gene expression dynamics (Fig. 1c). Considering the possibility that a similar interaction strength between LR pairs can yield diverse impacts on downstream gene expression dynamics^3^, it becomes crucial to assess the conversion rate for each receptor. Next, our approach involves the computation of the ligand diffusion rate (*γ*^*l*^ = *x*^*l*^/d^*τ*×*δ*^), which quantifies the ligands secreted by the sender cell that reach the target receiver cell (Fig. 1d; Eq 1 in Methods). This rate is a function of both ligand expression, *x*^*l*^ and intercellular distance, *d*. The ligand diffusion rate also governed by two parameters: *τ* and *δ*. *τ* represents the degrees of spatial freedom concerning the single-cell data, typically set to two for spatial transcriptomics data that is derived from two-dimensional slices. *δ* signifies the decay rate of ligand diffusion which prioritizes interactions over long or short distances. When *δ* is lower than 1, it weakens the decay rate, giving precedence to long-range interactions. Conversely, when *δ* exceeds 1, it amplifies the decay rate, favoring short-range interactions. A *δ* = 1 (default) equally prioritizes long and short-range interactions (no preference). Consequently, adjusting the *δ* parameter can aid in identifying LR pairs more inclined to interact over long-range or short-range distances. This could also identify proper *δ* rates for different ligands. We quantify the interaction between a specific LR pair from sender and receiver cell agents using LR scores, which depend on the above rates (Fig. 1d; Eq 4). The LR score for two cluster pairs is the Interaction Score (IS) (Fig. 1d; Eq 7). We consider an IS for an LR pair significant if its IS score significantly surpasses the background score derived from a permutation test. The cumulative interaction between two cell clusters is computed as the total number of significant LR pairs (Eq 8). CellAgentChat also computes the cell receiving score (CRS) (Eq 5) to measure the received interactions for each individual cell. Together, CellAgentChat offers a suite of downstream functionalities (Fig. 1e), several of which are distinctive and not available in existing methodologies.

### CellAgentChat identifies signaling pathways involved in mouse hippocampus development using high resolution spatial transcriptomics data

We first studied CCI in in mouse hippocampus development using time-series single-cell spatial transcriptomics data obtained from the Stereo-seq platform to accurately define the cellular transcriptomics and cellular composition in mouse neurogenesis^15^. Stereo-seq attains a nanoscale resolution of 500 nm, facilitating precise spatial pinpointing and characterization of individual cells^15^. We aimed to identify the key signaling pathways involved in the developmental lineage between cells at embryonic day E12.5, E14.5 and E16.5, which respectively contained 3,671 cells, 3,648 cells and 19,419 cells across 3 slices, from a total of 11 cell type populations (Fig. 2a).

**Figure 2:**
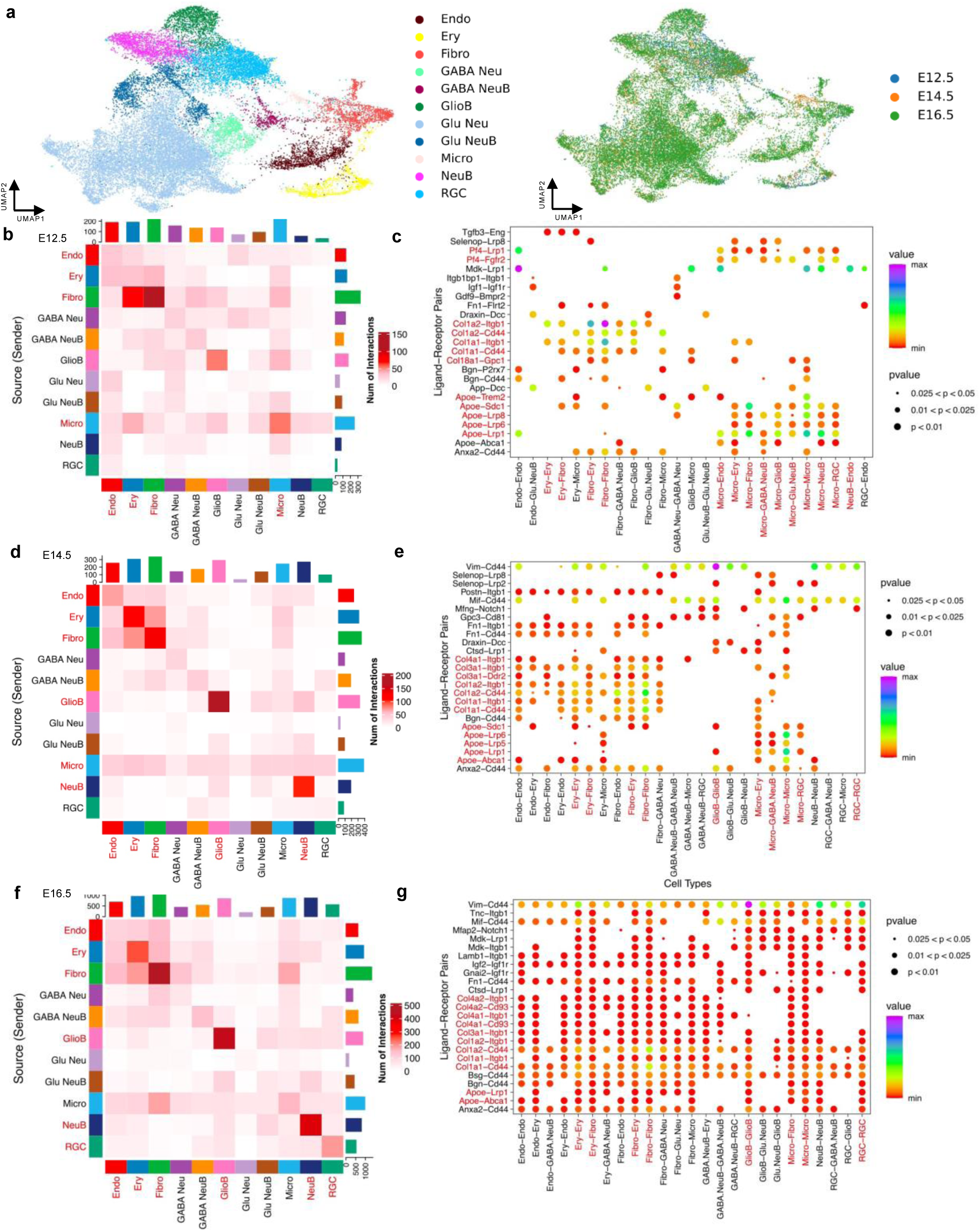
Analysis of cell communications in developing mouse hippocampus. **a**, UMAP visualization of scRNA-seq data across time points. (**b, d, f,)** Heatmaps display the communication network inferred by CellAgentChat at E12.5, E14.5 and E16.5, respectively. (**c, e, g,)** Dot plots illustrate the top 25 LR pairs between cell types inferred by CellAgentChat at E12.5, E14.5 and E16.5, respectively.

CellAgentChat calculates the interaction score based on the spatial coordinates of cells and the distance between individual cells when spatial information is provided (Fig. 1). When considering the spatial coordinates of cells, CellAgentChat identified 1,531 significant interactions involving 579 LR pairs at E12.5 (Fig. 2b, c). Microglial (Micro), Fibroblasts (Fibro), Erythrocytes (Ery), Endothelial cells (Endo) and Glioblasts (GlioB) were involved in the greatest number of interactions (Fig. 2b). The well-established description of Micro and other vascular cells’ role in promoting and regulating neurogenesis is widely acknowledged^30–32^. Micro also contributes to regulating the size of the neural precursor pool, which is crucial in the process of neurogenesis^30^. Upon examining the LR pairs with the highest involvement as inferred by CellAgentChat, it becomes apparent that the *Apoe* signaling pathway played a significant role (Fig. 2c). Interestingly, the top four LR pairs all centered around the *Apoe* ligand, with *Apoe-Lrp1* ranking at the forefront. The potential regulatory role of *Apoe* in mouse neurogenesis has been previously documented, including its involvement in Micro expression to mitigate neurodegenerative diseases^33,34^. Further, we see heavy collagen signaling (Fig. 2c). *Col1a1*, *Col1a2* and *Col8a1* are all commonly expressed by Fibro to enhance hippocampal neurogenesis in mice^35,36^. We also see *Pf4* signaling (Fig. 2c), which has also been shown to enhance hippocampal neurogenesis in mice^37^.

At E14.5 and E16.5, we observe a continuation of the similar trends seen at E12.5. There is consistently strong interaction among Micro and vascular cells such as Ery, Endo, and Fibro (Fig. 2d, f), highlighting their crucial role throughout neurogenesis^30–32^. Additionally, there remains significant *Apoe* signaling within Micro and collagen signaling originating from Fibro (Fig. 2e, g). However, notable changes emerge at E14.5. Firstly, there is increased signaling within glioblasts (GlioB) and neuroblast cells (NeuB) (Fig. 2d, e). Beginning at E14, radial glial cells (RGCs) produce intermediate progenitor cells like GlioB and NeuB before these cells terminally differentiate into specific neuronal types^38,39^. This phenomenon persists until E18, explaining the continued trend observed at E16.5 (Fig. 2f, g). Furthermore, there is an increase in interactions involving RGCs at E16.5 (Fig. 2f, g), coinciding with their transition from producing neurons to generating astrocytes and later oligodendrocytes from E16 until birth^38,39^.

The data presented in Fig. 2f illustrates the communication patterns at stage E16.5, combining three spatial sections into a cohesive analysis. We also analyzed the results inferred by CellAgentChat for each of the three slices at E16.5 individually (Supplementary Fig. 1). Across all three slices, we observe remarkably consistent communicative patterns, mirroring those seen in the combined analysis presented in Figure 2f. This consistency underscores the robustness of the tool, CellAgentChat. Within each of the three slices, robust signaling is evident among Micro, Ery, Endo, and Fibro cells, consistent with previous discussions (Supplementary Fig. 1a, b, c). Additionally, we observe strong correlations among these communication patterns across the slices (Supplementary Fig. 1d).

Since spatial information is not always provided, to showcases its broad applicability, we also applied CellAgentChat to the Stereo-seq mouse hippocampus development without using the spatial information. This led to consistent findings to those obtained from the spatial transcriptomics analysis including the crucial role of Micro and vascular cells and the increased levels of collagen and *Apoe* signaling to regulate neurogenesis (Supplementary Fig. 2).

To systematically assess the reliability of the LR interaction pairs predicted by CellAgentChat, we further employed Reactome pathway enrichment analysis. We compared the enrichment P-values of these pathways’ terms using the significant LR pairs with high scores (**Methods**). Many pathways associated with the LRs predicted by CellAgentChat are related to neurogenesis and are present in all time points and scenarios (spatial and non-spatial). These pathways included axon guidance, nervous system development and developmental biology. One of the most significant Reactome pathway in all time points was extracellular matrix (ECM) organization (Supplementary Fig. 3). Additional pathways involving ECM interactions had high adjusted P-values, indicating that these interactions play a role in neurogenesis (Supplementary Fig. 3). Studies have shown that the ECM influences and regulates many aspects of neurogenesis in mouse and other species^40,41^. Pathways involving collagen were also present (Supplementary Fig. 3), including collagen degradation which supports the collagen signaling discussed earlier. Collagen is the most abundant component in the ECM and is broken down with the help of Micro cells expressing *Mmp9* to enhance neurogenesis^42^. Finally, we see *EPH-Ephrin* signaling (Supplementary Fig. 3), which regulates cell migrating neurons in nervous system development^43^.

Together, we showcased the utility of CellAgentChat for inferring CCIs with high-resolution spatial transcriptomics data by recapitulating the key time-dependent signaling events during the developmental process of neurogenesis in mice. Moreover, its consistent performance across various samples, including different time-points and technical variations, speaks to its reliability and robustness. Additionally, CellAgentChat can seamlessly integrate technical samples, such as different spatial slices, into a unified analysis framework, thus facilitating a comprehensive understanding of complex cellular interactions and signaling dynamics in spatially resolved datasets. CellAgentChat demonstrates versatility in dealing with scRNA-seq datasets with and without spatial information.

### CellAgentChat finds signaling pathways in spatial transcriptomics data at multi-cell/spot resolution

To showcase the broad applicability of CellAgentChat, we further explored its utility in signaling analysis using Visium spatial transcriptomics data, where each spatial spot contains multiple cells. This approach aimed to demonstrate that CellAgentChat can efficiently operate at the spot-level in single-cell spatial datasets, which remain commonly adopted for single-cell spatial studies. In this application, we term spots as agents in the ABM. We applied CellAgentChat to a dataset of Squamous Cell Carcinomas (SCC), which comprises 3,295 spots from a total of thirteen distinct cell populations, corresponding to three normal keratinocyte (NKer) clusters, two tumor keratinocyte (TKer) clusters, and three myeloid dendritic cell (MDC) clusters (Fig. 3a). Additionally, the dataset includes several other cell types, such as endothelial (Endo), fibroblast (Fib), T-cells, B-cells, and melanocytes (Mel). When considering the spatial coordinates of spots, CellAgentChat identified 7,973 significant interactions involving 1521 LR pairs (Fig. 3b, c). Only a few cell types were involved in these interactions. NKer3, MDC3, T cells, and TKer2 participated in the most interactions (Fig. 3b). Furthermore, CellAgentChat identified that three out of the four LR pairs engaged in the most interactions prominently featured *CD44* as the receptor, with *PKM-CD44* emerging as the top pair (Fig. 3c). This discovery aligns with previous studies demonstrating the pivotal role of this LR interaction in the progression of SCC^44^. Additionally, the *MMP1-CD44* pair was involved in numerous interactions, with *MMP1* highly expressed in TKer subpopulations and *CD44* overexpressed in cancer cells (Fig. 3c). These findings align with previous studies linking *MMP1* expression to SCC, highlighting the significance of *CD44* in cancer cells^45,46^. The LR pair *HLA-B-CANX* also stood out as a leading LR interaction, notably elevated in T cells and TKer2 cells (Fig. 3c). This compelling association has been demonstrated to be linked with SCC^47–49^.

**Figure 3:**
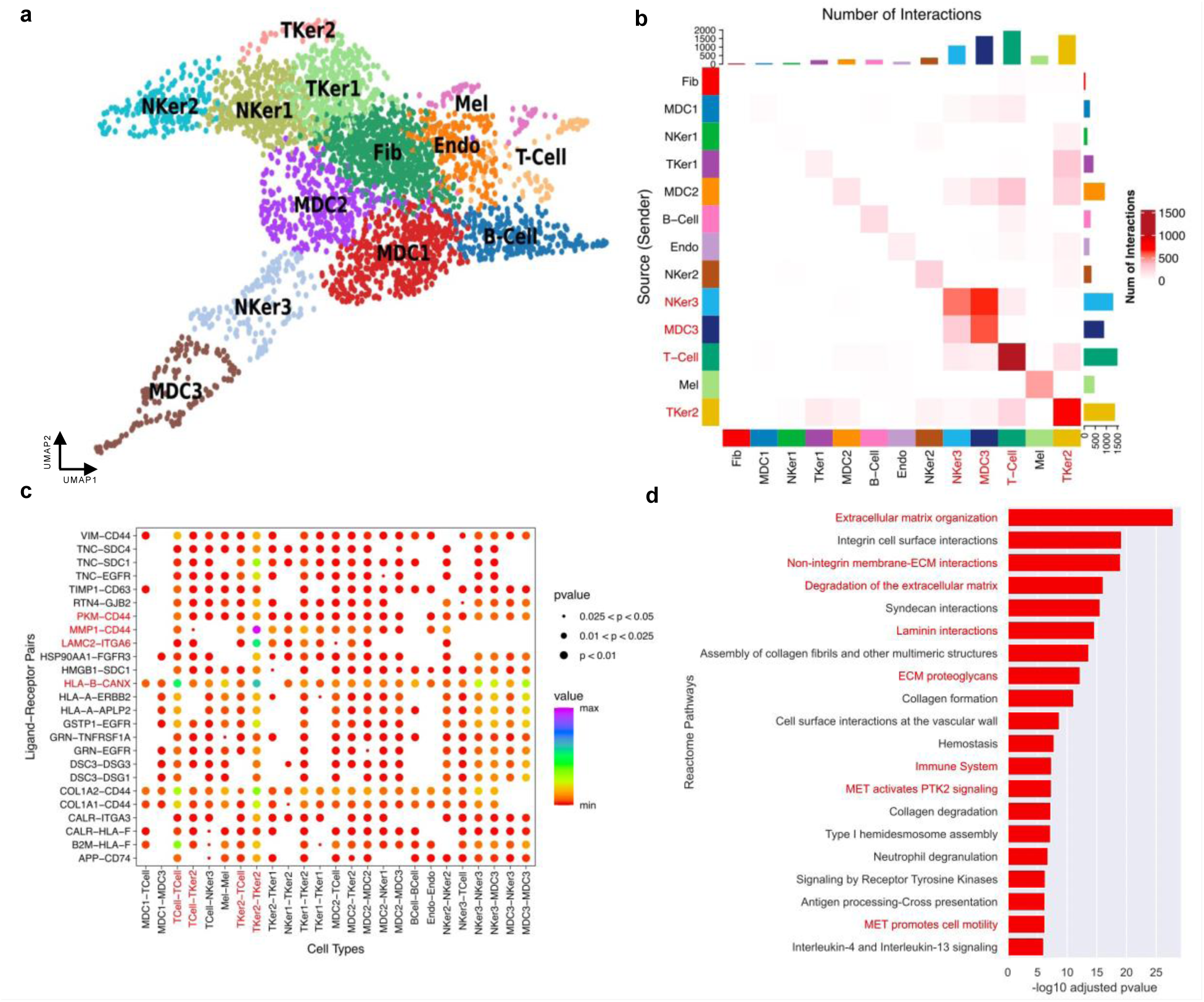
Analysis of Cell-Cell Communications in Squamous Cell Carcinoma (SCC) using CellAgentChat. **a**, UMAP visualization of spot resolution single-cell transcriptomics data across time points. **b,** Heatmap displays the communication network between cell types, inferred by CellAgentChat. **c,** Dot plot illustrates the top 25 LR pairs involved in the communication between cell types inferred by CellAgentChat. **d,** Reactome pathways analysis using the ligand-receptor interactions inferred by CellAgentChat. Pathways highlighted in red are linked with SCC.

Additionally, we employed Reactome pathway enrichment analysis to assess the reliability of the LR interaction pairs predicted by CellAgentChat. Many pathways associated with the LRs predicted by CellAgentChat are relevant to the disease of interest. The most significant Reactome pathway was ECM organization (Fig. 3d). Additional pathways involving ECM interactions had high -log_10_ adjusted P-values, indicating that these interactions play a role in SCC. Studies have shown that the ECM regulates cell behavior and tumor progression in SCC^50^. The immune system pathway was also observed (Fig. 3d). Cancer patients often experience an elevated immune response, with T-cells playing a critical role in recognizing and responding to tumor antigens in all types of cancers, including SCC^51^. MET signaling pathways were also discovered (Fig. 3d). MET signaling pathway regulates a range of downstream processes, including proliferation and invasiveness of tumor cells, and researchers have associated it with poor prognosis in SCC^52,53^. Lastly, interaction pathways involving Laminin were discovered (Fig. 3d). Laminins constitute integral components of ECM basement membranes, directly governing essential biological processes linked to morphogenesis, including proliferation, adhesion, migration, angiogenesis, and cell survival^54^. Deviations in laminin expression patterns and functionality are implicated in pathological conditions, such as carcinogenesis, due to their ability to modulate critical cellular processes, thereby influencing various cellular behaviors^54–56^. Furthermore, numerous studies have indicated that alterations in laminin expression patterns and activity within tumor tissues, including SCC, correlate with patient outcomes, such as tumor invasiveness and prognosis^54–56^. The inclusion of this pathway is supported by the identification of *LAMC2-ITGB6* as a key LR interaction involving TKer2 cells (Fig. 3c).

### Effective in silico receptor blocking by CellAgentChat for analyzing perturbed downstream genes and potential therapeutics

We harnessed CellAgentChat to build upon previous research on breast cancer exploring the Xenium human breast cancer dataset^16^, aiming to identify significant receptors related to the disease and cultivate innovative therapeutic approaches. The first step of our process entailed using CellAgentChat to pinpoint 13 key receptors involved in the top 25 LR interactions for the cellular communications in the dataset (Supplementary Fig. 4d, 5a). We then simulated the blocking of these receptors individually, akin to the actions of receptor inhibitor drugs. We deployed a feedforward neural network that was designed to calculate each receptor’s conversion rate and was trained to predict the expression of all genes using the ligand and receptor expression as inputs. The network connections were constrained to the known LR interactions, LR-TF, and TF-genes along the forward pass, respectively. (Fig. 4a). Using the trained network, we predicted the potential impact of receptor blocking on downstream gene expression and identified the most perturbed downstream genes for each in silico blocked receptor. We subsequently assessed these perturbed genes using DisGeNET^57^ enrichment analysis to evaluate the potential therapeutic implications of blocking specific receptors (Fig. 4a). Our analysis with CellAgentChat indicated that blocking 9 of the 13 identified receptors—specifically EGFR, PDCD1, PECAM1, SDC4, CXCR4, AVPR1A, CTLA4, CD80, and CD93 (hereafter referred to as candidate receptors)— significantly disrupts genes associated with breast cancer as listed in DisGeNET (Fig. 4b). Further investigation showed that the blockade of four specific receptors—PECAM1, CXCR4, CTLA4, and CD80—also resulted in substantial expression changes of downstream genes that significantly overlapped with breast carcinoma target genes in DisGeNET (binomial test; *P* < 0.05)

**Figure 4:**
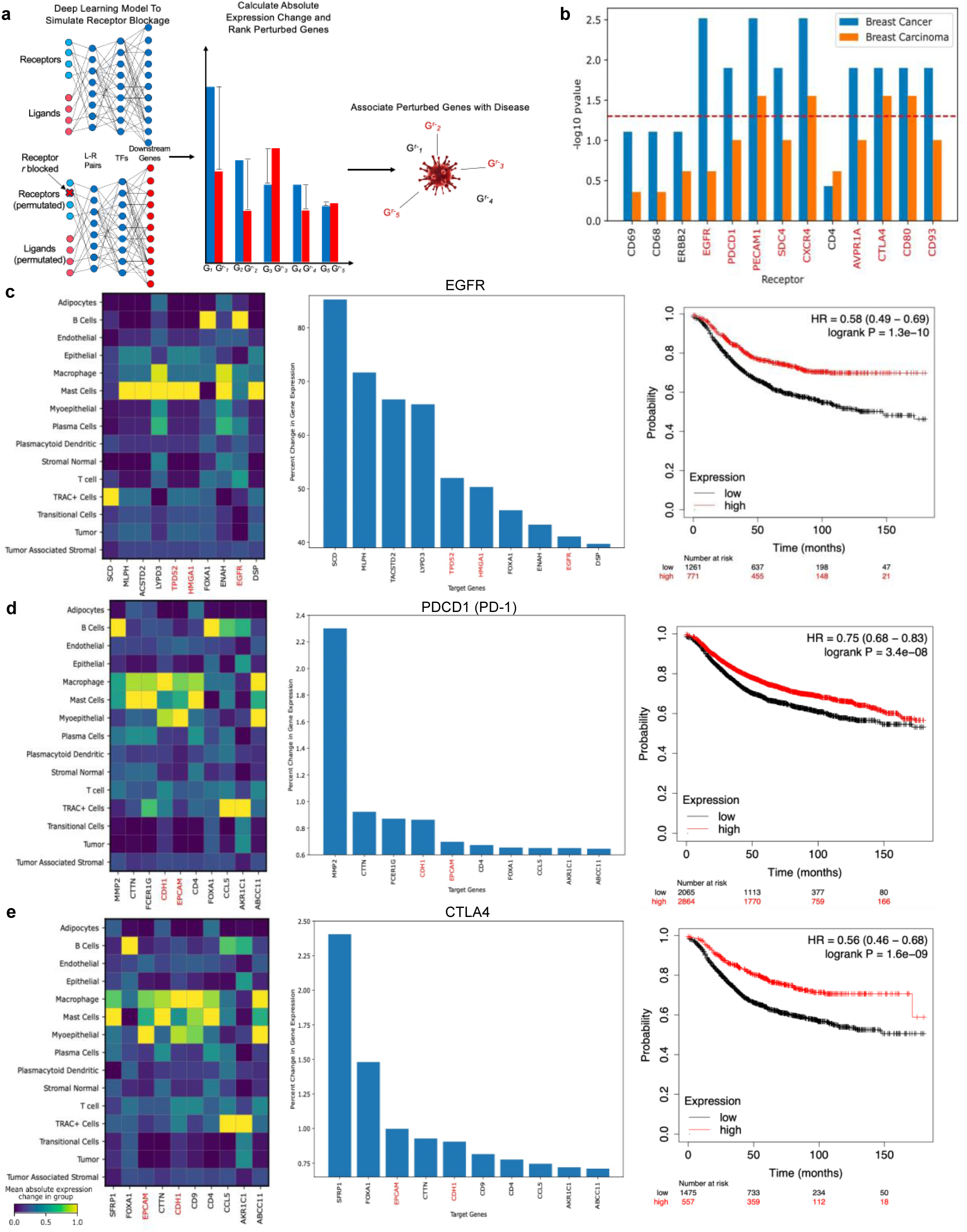
Identification of key downstream genes related to breast cancer after in silico blockage of top interacting receptors. **a**, Schema of the in-silico perturbation analysis using our neural network to block receptors and identify the most perturbed downstream genes linked with breast cancer. **b,** Enrichment for the breast cancer and breast carcinoma gene sets of the significantly perturbed genes based on the in-silico perturbation of the selected 13 receptors. The enrichment is represented as the -log_10_ adjusted P-values (binomial test; FDR corrected-Bonferroni Correction). *EGFR*, *PDCD1*, *PECAM1*, *SDC4*, *CXCR4*, *AVPR1A*, *CTLA4*, *CD80* and *CD93*, highlighted in red, have significant -log_10_ adjusted P-values for the breast cancer gene set enrichment. (**c, d, e,)** Matrix plots display the mean change in expression of each target gene for each cell type for, *EGFR*, *PDCD1* (*PD-1*) and *CTLA4*, respectively (left). Identification of the top 50 downstream genes with the highest percent change in absolute expression upon blocking *EGFR*, *PDCD1* (*PD-1*) and *CTLA4*, respectively (middle). A red highlight indicates genes associated with breast cancer. Survival analysis displaying the probability of survival of high vs. low receptor expression groups for *EGFR*, *PDCD1* (*PD-1*) and *CTLA4*, respectively (right).

In an ablation study to highlight the benefits of integrating biological priors, we tested a fully connected feedforward network without the LR-TF and TF-target prior constraints that were used to define connectivity in our primary analysis. This approach aimed to demonstrate the impact of our initial incorporation of biological insights. The removal of these biological constraints resulted in a noticeable decrease in specificity: four receptors previously identified as nonsignificant were now falsely marked as significant (Supplementary Fig. 5b).

The cell receiving score (CRS) of each cell, as identified from our animation platform, reveals that tumor cells, tumor-associated stromal cells, endothelial cells, and macrophages demonstrate the highest degrees of involvement in interactions with the candidate receptors (Supplementary Fig. 6). We have also observed significant interactions among these populations, bolstering this conclusion (Supplementary Fig. 4c). The substantial engagement of tumorous and immune cells in interactions with the candidate receptors further validates the precision of our cellular interaction analysis and underscores the potential effectiveness of receptor inhibitor drugs that target these critical cellular interactions. Following the blocking of receptors, interactions among these cell types will be entirely inhibited, resulting in a significant alteration of the cells’ interaction profile.

Although studies have associated all of the candidate receptors examined in this study with breast cancer, the candidate receptors we identified—*EGFR*, *PDCD1* (*PD-1*), and *CTLA4*—appear to play a particularly crucial role in breast cancer development^58–60^ (Fig. 4c-e). Furthermore, therapeutic treatments targeting the three receptors in breast cancer have been previously documented. Both Lapatinib and Pembrolizumab are FDA approved drugs designed to specifically target and disrupt pathways involving *EGFR* and *PD-1*. Ipilimumab and tremelimumab are approved by the FDA to suppress the role of *CTLA4* in melanoma^60^. Although ipilimumab and tremelimumab are not FDA-approved for breast cancer treatment, studies have shown their positive impact in this context^60^. This corroborates the potential application of CellAgentChat to identify receptor-mediated therapeutics. Moreover, blocking the *EGFR*, *PDCD1* (*PD-1*), and *CTLA4* receptors in silico perturbs the disease genes associated with breast cancer (from DisGeNET) (Fig. 4c-e). In particular, when *EGFR* was blocked, *TPD52* emerged as the fifth most affected gene, previously linked with breast cancer^61^ (Fig. 4c). Similarly, *CDH1*, another gene associated with breast cancer^62^, was the fourth most affected gene when *PDCD1* (*PD-1*) was blocked (Fig. 4d), and *EPCAM*^63^ was the third most affected gene when *CTLA4* was blocked (Fig. 4e). Numerous perturbed genes notably influenced immune cell types like mast cells and macrophages (Fig. 4c-e; left). This observation aligns with the context of breast cancer, where immune cells play a crucial role in the tumor microenvironment and the immune response against cancer cells. All other breast cancer disease target genes perturbed by blocking the candidate receptors are highlighted in red in (Fig. 4c-e, Supplementary Fig. 5c-e, 7a-c).

Finally, to validate the accuracy of the candidate drug targets captured by CellAgentChat, we conducted a survival analysis of breast cancer patients over time, comparing populations with high and low expression levels of each candidate receptor. The analysis revealed statistically significant differences in survival probabilities over 150 months for patients with varying levels of expression of *EGFR*, *PDCD1* (*PD-1*), and *CTLA4* (Kaplan-Meier log-rank test; *P* = 1.3x10^-10^, *P* = 3.4x10^-8^, *P* = 1.6x10^-9^, respectively) (Fig. 4c-e). Additionally, survival analysis of all other candidate receptors exhibited statistically significant differences in survival probabilities between the two patient groups (Supplementary Fig. 5c-e, 7a-c). These findings indicate that the identified candidate receptors would serve as suitable therapeutic targets for treating breast cancer.

In addition to using CellAgentChat to perform in-silico perturbation analysis on high-resolution spatial transcriptomics data of a human breast cancer dataset, we also demonstrate the in-silico perturbation capabilities of our model on a spot-level resolution pancreatic ductal adenocarcinoma (PDAC) dataset, where we identified *NOTCH3*, *PLAUR*, *ITGA11* and *CD74* as potential therapeutic targets (see Supplementary Text; Supplementary Figs. 8 and 9).

### CellAgentChat facilitates the agent-based visualization of cellular interactions associated with individual cells

We employed an ABM approach through CellAgentChat, allowing us to examine individual cell interactions rather than solely focusing on cell populations. We applied the method to a spot-level human pancreatic ductal adenocarcinoma (PDAC) dataset^64^. After preprocessing the PDAC dataset, we discerned six distinct cell populations: ductal (Duct), three subpopulations of acinar (Acin1, Acin2, and Acin3), stromal (Strom) cells, and two classes of cancer (Can1 and Can2) (Supplementary Fig. 4b, e, f). To effectively exhibit the insights that CellAgentChat can provide into individual cell interactions, we identified twenty most receiving spots (group A, red circles) based on the cell receiving score (CRS) (**Methods**) and the least receiving spots (group B, black circles) with the lowest CRS within the second cancer cell class (green cells) (Fig. 5a).

**Figure 5:**
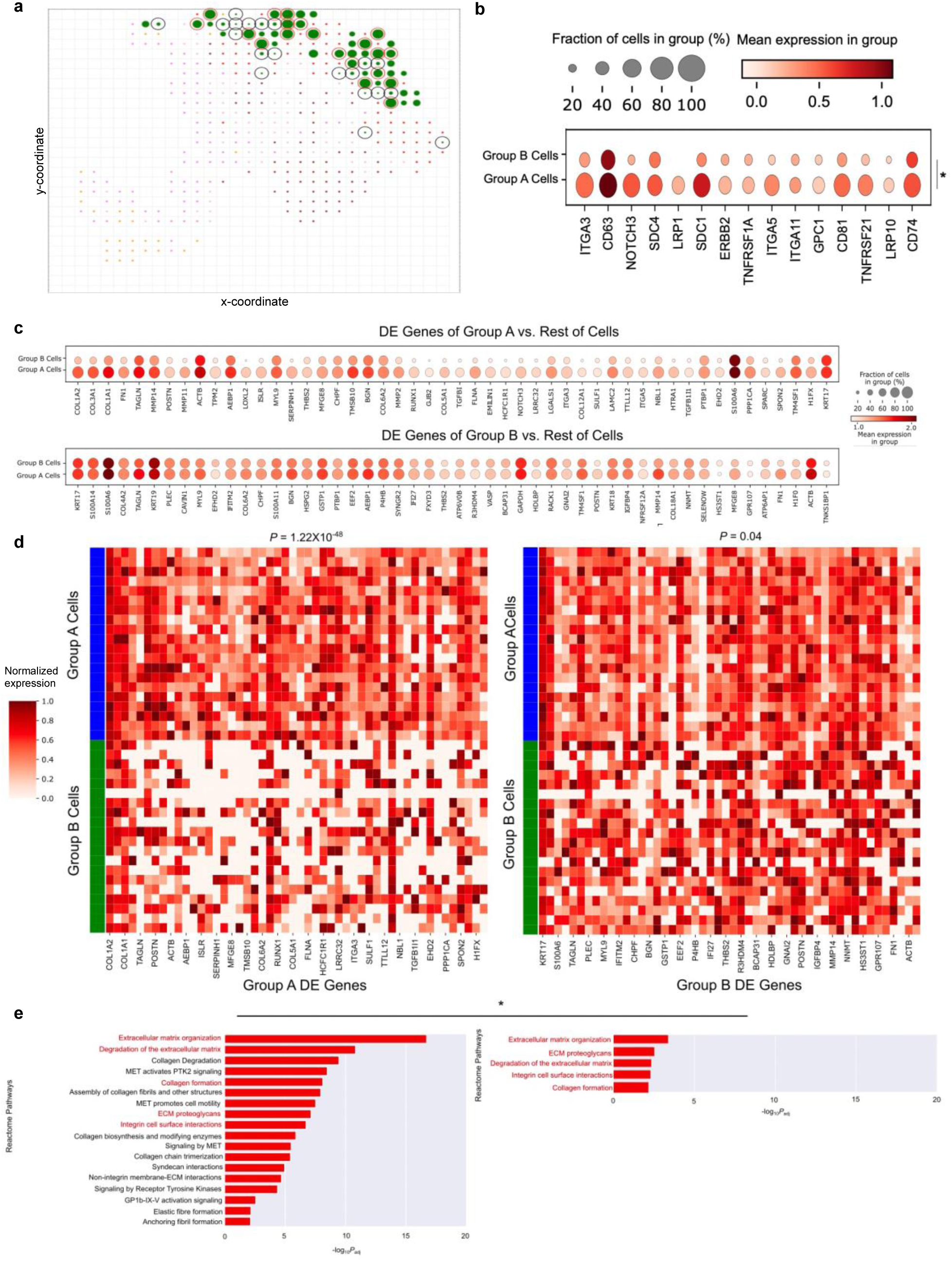
Application of CellAgentChat to detect variations in intercellular communication among individual cancer cells in PDAC. **a**, CellAgentChat’s animation platform enables the detection of communication strength variations in individual cancer cells. The platform highlights the cell receiving score (CRS) (**Methods**) of cancer subpopulation one cells (colored green). It identifies the top 20 cells with the highest CRS (group A) marked by red circles and the bottom 20 cells with the lowest CRS (group B) marked by black circles. **b,** Expression analysis of receptors involved in the top 25 ligand-receptor interactions reveals that Group A, consisting of the top 20 spots with the highest CRS identified by CellAgentChat’s animation platform, shows significantly higher expression of key receptors compared to Group B, consisting of the bottom 20 receiving cancer spots (Wilcoxon test one-sided; *P* < 0.05). **c,** Mean expression of differentially expressed (DE) genes up-regulated for group A spots relative to rest of the spots (top) and up-regulated for group B cells relative to rest of the spots (bottom). **d,** Heatmaps display the differentially expressed (DE) genes for group A (left) and group B (right). The DE genes of group A cells are mainly expressed by group A and not by group B (Wilcoxon test one-sided; *P* < 0.001). Both groups express the DE genes of group B but are more significantly expressed by group A spots (Wilcoxon test one-sided; *P* < 0.05). **e,** The results of the Reactome pathway analysis using group A DE genes (on the left) and group B DE genes (on the right). Pathways affected by PDAC are more associated with group A DE genes (Wilcoxon test one-sided; *P* = 0.0313). Red highlights indicate shared pathways.

The spots with higher CRS (more interactive) generally reside in the outer layer of the tissue (invasive front), consistent with the discovery from many existing studies^65^. These two cell groups of distinct interacting patterns within the same cluster also demonstrated significant transcriptomic differences. Group A showed higher mean expression levels of receptors involved in the top 25 LR pairs than Group B (Fig. 5b). For example, *CD63* was highly expressed in both groups, with group A having the highest expression levels (Fig. 5b). Additionally, receptors such as *LRP1*, *SDC1*, *ITGA3*, *NOTCH3*, and *CD74* displayed notable differences in mean expression between the two groups, with group A again exhibiting higher expression (Fig. 5b). Next, we systematically explored the differentially expressed (DE) genes of these two groups. To identify the DE genes for each group (A and B), we compare the spots within the group with the rest of the spots in the dataset. The differential genes associated with group A show significantly higher expression in group A compared with group B (Wilcoxon one-sided test; *P* =1.22x10^-48^) (Fig. 5c). In contrast, the expression of group B DE genes is relatively similar across groups (Fig. 5c).

We showed the top 50 DE genes for both groups by comparing the expression of spots in groups A and B (Fig. 5d). While both groups showed high expression levels of *S100A6* and *ACTB*, group B cells exhibited elevated expression of *KRT19* (Fig. 5d). Conversely, *COL1A1*, responsible for type I collagen production, was particularly prominent in group A cells (Fig. 5d). Notably, *COL1A1*, a gene found enriched in PDAC tumor cells, is associated with tumor growth and metastasis in cells^66^.

To identify the biological functions associated with these groups, we conducted Reactome pathway enrichment on the top 50 DE genes for both groups (Fig. 5e). Intriguingly, group A was associated with several pathways related to the extracellular matrix (ECM) and collagen, known to be crucial in PDAC tumor progression^67^. On the other hand, group B showed less involvement in ECM-related pathways, aligning with their lower level of interaction compared to group A (Fig. 5e). Examinations of CCIs revealed that cells of the same cluster/type can display diverse interaction behaviors and play varying roles in pathogenesis and disease progression.

### CellAgentChat can be tuned to identify long and short-range interactions

Our understanding of the specific distances at which LR pairs operate is evolving but remains incomplete. While some interactions occur closely between cells, others span longer distances, influenced by factors such as specific ligands and receptors, tissue environment, and physiological conditions. The ABM employed by CellAgentChat provides nuanced representations of cell agent behaviors, including the diffusion of signaling ligands. As a result, this facilitates modeling of both short- and long-range interactions at the single-cell level by adjusting the delta (δ) factor to manipulate the ligand diffusion rates (**Methods**).

Using CellAgentChat, we embarked on an investigation on the spatial distance tendencies of cellular communication within the mouse somatosensory cortex leveraging data from SeqFISH+ technology (Supplementary Fig. 10). To this end, we utilized the CellChat database, which provides known annotations of the distance ranges for each ligand-receptor (LR) pair. LR pairs are categorized as either "Secreted Signaling" (indicating long-range interactions) or "cell-cell contact" (indicating short-range interactions). To identify short-range interactions, we used a δ=10, to constrain interactions to shorter distances and used δ=0.1 to identify long-range interactions. To validate our results, we compared the top interacting LR pairs with CellChat’s annotations. Under our short-range mode, the majority of our top 10, 15, 20 and 25 LR pairs were annotated as “cell-cell contact” (short-range) (Fig. 6a). Specifically, 80% of the pairs within the top 15 LR pairs identified in the short-range mode fell into the "cell-cell contact" category (Fig. 6a). In the long-range mode, most of the top LR pairs identified were classified as "secreted signaling" (long-range), with 75% of pairs within the top 15 LR pairs falling into this category (Fig. 6a). Recently, Liu et al.^68^ have developed a method to identify long-range and short-range interactions using optimal transport (OT). Compared to that approach, CellAgentChat more accurately captured short-range interactions and identified long-range interactions to a similar degree of accuracy (Fig. 6b).

**Figure 6:**
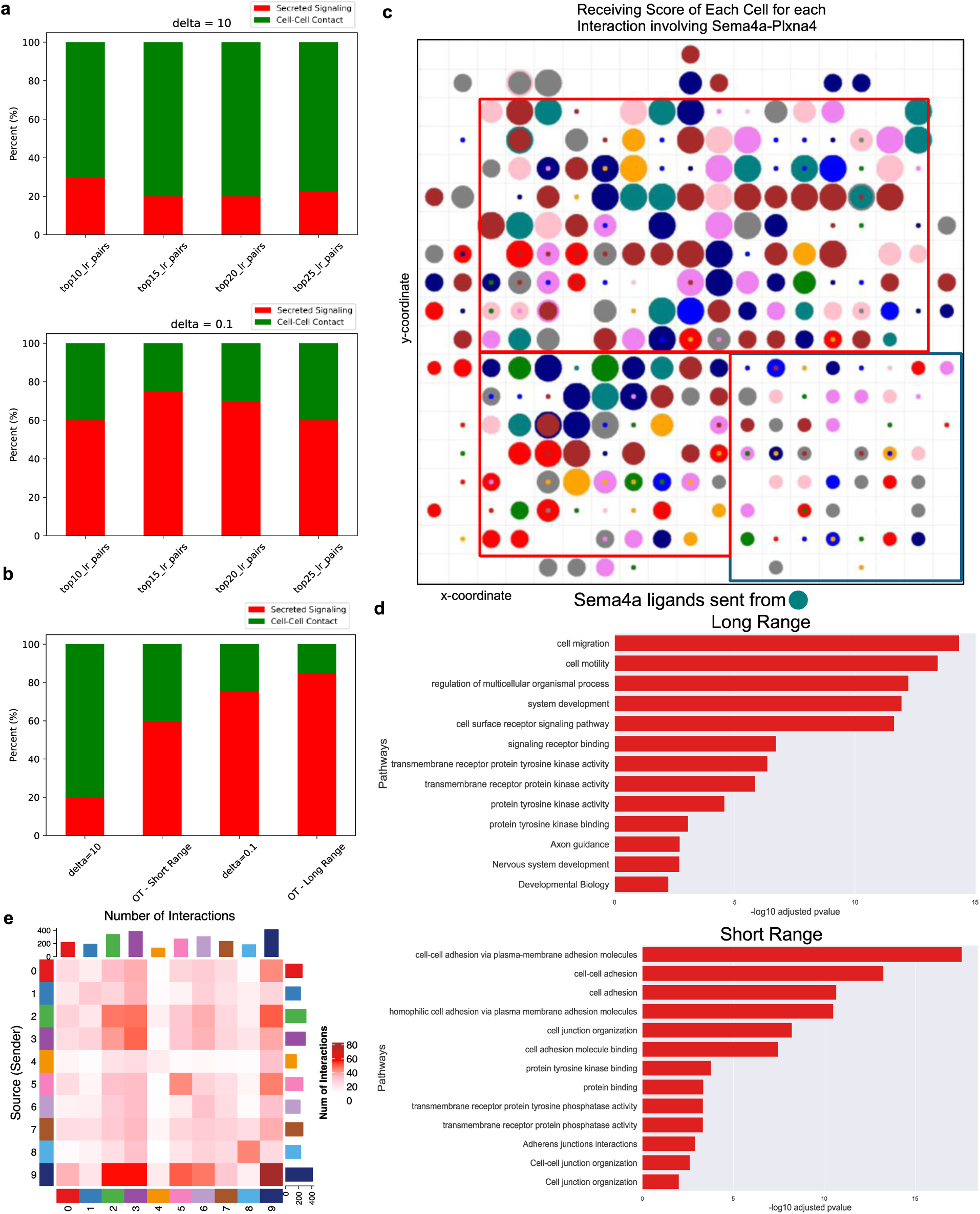
Application of CellAgentChat to infer long-range and short-range interactions in adult mouse cortex. **a**, Bar plot illustrates the percentage of interactions found by CellAgentChat annotated by CellChat’s database as either cell-cell contact or secreted signaling with a delta (δ) value set to 10 (top) and 0.1 (bottom). **b,** Bar plot illustrates the percentage of interactions found by CellAgentChat annotated by CellChat’s database as either cell-cell contact or secreted signaling in comparison to an optimal transport-based approach^68^. **c,** CellAgentChat’s animation platform depicts the cell receiving score (CRS) for the *Sema4a-Plxna4* ligand-receptor pair using the short-range mode (δ =10). The colour of the cells (circles) represents the cell cluster. This animation depicts the release of *Sema4a* (ligand) from cells in cluster 9 (represented by turquoise cells). In the animation, cell size corresponds to the cell receiving score (CRS). Cells closer to cluster 9 cells exhibit higher CRS (within red square) than cells that are further away (within blue square), which are smaller in size. **d,** Reactome pathway analysis of the receptors involved in the top 25 LR interactions in the long-range mode (top) and short-range mode (bottom). **e,** Heatmap displaying the cell communication between clusters after prescribing specific delta values based on identifying key short-range and long-range interactions.

In our short-range mode, we detected LR pairs involving the *Sema4* pathway, such as *Sema4a- Plxna4*. Previous research has linked these pairs to axon guidance in the mouse cortex, occurring over short distances^69^. We further employed our animation platform to visualize the CRS of individual cells for the *Sema4a-Plxna4* LR pair in the short-range mode originating from cluster 9 cells (turquoise cells) (Fig. 6c). The results highlighted higher CRS amongst cells close in proximity to the cluster 9 cells represented by larger sizes (within the red box), compared to cells situated further away (blue box). This observation underscores the capabilities of CellAgentChat in modeling and identifying short-range interactions.

We conducted pathway enrichment comparisons of the LR interaction pairs predicted by CellAgentChat for both long- and short-range interactions (Fig. 6d). The genes associated with short-range interactions showed enrichment in pathways related to cell adhesion and cell-cell junctions. In contrast, genes associated with long-range interactions were enriched in numerous signaling pathways known for their wide regulatory range (Fig. 6d).

Ultimately, we leverage our understanding of the above discovered short-range and long-range LR pairs to refine the ligand diffusion rate with greater precision. Having pinpointed distinct LR pairs engaging in either long-range or short-range interactions, we assign specific delta values to these ligands, to control the decay rate between cell *i* and *j* in computing the ligand diffusion rate *λ*^*l*^ (Eqn. 1). This enables us to more accurately depict and simulate their actual decay rates during the computation of CCIs. By integrating the recommended decay rates for the most impactful long and short-acting ligands, we detect both long-range and short-range interactions concurrently (Fig. 6e). This reveals a communication pattern resembling interactions observed solely with scRNA-seq data (Fig. 6e, Supplementary Fig. 10b), where spatial constraints are absent, thus more accurately facilitating long-range interactions. Nevertheless, the communication pattern also continues to exhibit numerous short-range interactions between proximate cells (e.g., cluster 9- cluster 9 cells, cluster 3-cluster 3 cells) (Fig. 6e).

### CellAgentChat empowers dynamic simulations using agent-based modeling

CellAgentChat provides dynamic simulation capabilities to analyze the impact of CCIs on cell states over short durations. At each time point (time step), CCIs are computed as described earlier, followed by the prediction of gene expression levels for each cell using our trained neural network (**Methods**). Due to uncertainties and the influence of various factors that could affect the actual magnitude of gene expression changes, directly substituting existing gene expression values with predicted ones is impractical. Instead, we opt for assessing the directional change of each gene— whether they increase or decrease—which proves to be more robust and easier to predict^70^ (Supplementary Fig. 11a). Genes with significant changes are identified for each cell at each time step, and their expression is adjusted within the ABM for the next time step (**Methods**) (Supplementary Fig. 11a). This iterative process is repeated across all time steps, reflecting the dynamic modulation of gene expression by CCIs.

To validate the accuracy of our dynamic simulations, we determined the correlation between the pseudotime trajectory of each gene, and its trajectory as established by the dynamic simulation of CellAgentChat. Across all genes, we observed a correlation between the pseudotime trajectory and the ABM dynamic trajectory (Supplementary Fig. 11b). Additionally, we examined the trajectory of highly interacting ligands and receptors, finding strong agreement between the pseudotime trajectory and the dynamic ABM trajectory for genes like *ITGA6* (receptor) and *MMP1* (ligand), which showed overall increasing trends over time (r^2^ = 0.93 and r^2^ = 0.97, respectively) (Supplementary Fig. 11c, d). Similarly, both trajectories indicated a decreasing expression trend for *CALR* (ligand) over time (r^2^ = 0.90) (Supplementary Fig. 11e). Moreover, CellAgentChat accurately identified the initial decrease of expression of *CD44* (receptor) followed by a rapid increase (r^2^ = 0.89) (Supplementary Fig. 11f). These findings highlight the effectiveness of CellAgentChat in capturing dynamic cellular changes resulting from CCIs within a time frame comparable to the pseudotime represented by the cells.

### Benchmarking of CellAgentChat with other cell–cell communication inference tools

We undertook a comparative analysis of CellAgentChat against five prominent CCI inference methods based on both scRNA-seq and spatially resolved transcriptomics data: CellPhoneDB (v5)^17^, CellChat (v2)^18,23^, NICHES^22^, COMMOT^21^ and Scriabin^24^. In our comprehensive evaluation of methods, we benchmarked them rigorously from several key perspectives. First, due to the absence of ground truth, we compared the agreement of the cell-cell communication networks between each benchmarked method with the consensus of all methods. Secondly, we examined the consistency of CellAgentChat’s predictions with spatial information, focusing on the spatial proximity of stronger interactions. Specifically, we measured the Pearson correlation between the probability of interaction and the inverse distance between cells, grounding this analysis on the established understanding that cells are more likely to interact when they are in closer proximity^3^. Thirdly, we assessed the functional relevance of the predicted LR pairs by conducting Reactome pathway enrichment analysis. Finally, we measured the overlap of inferred LR pairs across different methods.

We began by benchmarking across methods using only scRNA-seq datasets to validate the robustness of our CCI calculation framework before integrating spatial data. We assessed the cell-cell communication networks generated by CellAgentChat and compared them with the outcomes of other methods. Due to the absence of a definitive ground-truth, we adopted a consensus/ensemble approach. This involved normalizing the interaction matrices computed by each method and averaging them to create an ensemble interaction matrix, serving as our new consensus reference. Subsequently, we computed the Pearson correlation between the ensemble interaction matrix and the inferred communication patterns of each method for comparison. Across all datasets, the results from CellAgentChat demonstrated superiority and consistency with the communication networks inferred by CellPhoneDB, CellChat, and NICHES, showing a higher median Pearson Correlation than the other three methods (Fig. 7a (top)). On examining individual datasets, CellAgentChat consistently outperformed CellPhoneDB and Scriabin in all cases (Wilcoxon one-sided test; *P* < 0.05) (Supplementary Fig. 12, 13, 14). Moreover, CellAgentChat, CellChat, and NICHES showcased remarkably consistent and high-performance results across all evaluated metrics (Supplementary Fig. 12, 13, 14). Upon incorporating spatially resolved transcriptomics data, CellAgentChat continued to perform with a high degree of consistency among the state-of-the-art methods. Specifically, CellAgentChat maintained superior performance compared to CellPhoneDB and also outperformed CellChat (Wilcoxon one-sided test; *P* < 0.05) (Fig. 7a (bottom), Supplementary Fig. 12, 15, 16). In comparison to NICHES and COMMOT, CellAgentChat performed comparably, further demonstrating its status among the state-of-the-art methods.

**Figure 7:**
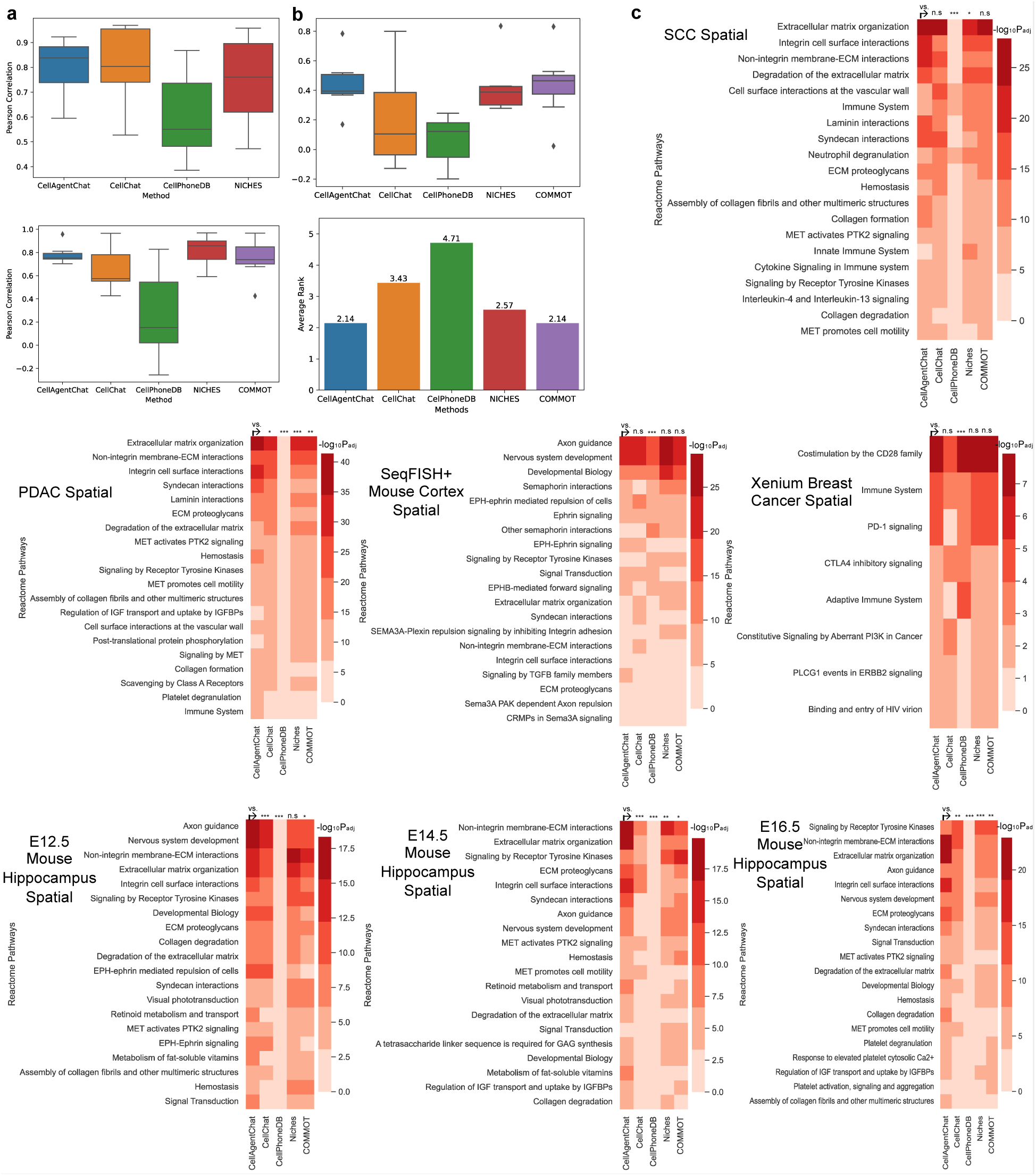
Benchmarking of CellAgentChat with existing state-of-the-art methods. **a**, Boxplot comparing the Pearson correlation between inferred cell communication by individual methods and the ensemble of all methods, for both non-spatial (top) and spatial data (bottom). **b,** Boxplot comparing the Pearson correlation between the inferred cell communication for each method and the inverse cell-distance matrix (top), across datasets. Bottom plot in panel shows the average rank across datasets. **c,** The Reactome pathway analysis, performed on receptors sourced from the top 100 ligand-receptor pairs inferred by each method across various datasets using spatial transcriptomics data, revealed that the pairs identified by CellAgentChat exhibit a greater level of statistical significance.

To validate that the CCIs inferred by CellAgentChat are spatially informed, we computed the Pearson correlation between the predicted communication matrix and the cell-cell inverse distance matrix. Indeed, cells predicted by CellAgentChat as highly interacting were significantly closer in proximity compared to predictions from other methods (Fig. 7b). We demonstrated that our spatially informed predictions outperformed those of CellPhoneDB and CellChat (Wilcoxon one-sided test; *P* < 0.05) (Fig. 7b), showing superior performance across every dataset compared to CellPhoneDB and in all but one dataset compared to CellChat (Supplementary Fig. 17). Furthermore, we show comparable performance compared to NICHES and COMMOT (Fig. 7b, Supplementary Fig. 17). Overall, CellAgentChat and COMMOT had the joint highest average rank across all datasets (Fig. 7b (bottom)).

After validating the accuracy of CellAgentChat in predicting cell-cell interactions (CCIs) through consensus and spatial proximity metrics, we proceeded to conduct pathway enrichment analysis to further confirm the biological relevance of the inferred LR pairs. This step serves as an additional layer of validation. We conducted computational validations, as previously described, comparing the enrichment P-values for relevant pathways using ligands and receptors from the top 100 significant LR pairs from each method. The pathway enrichment analysis conducted for the results using only scRNA-seq data underscored CellAgentChat’s superiority over other methods. CellAgentChat performed better than CellPhoneDB and Scriabin across all datasets (Supplementary Fig. 18). Additionally, CellAgentChat outperformed CellChat in five out of the seven datasets and showed comparable performance in the remaining two (Supplementary Fig. 18). When compared to NICHES, CellAgentChat performed better on the PDAC, breast cancer, and E16.5 mouse hippocampus datasets, while demonstrating comparable performance on the rest (Supplementary Fig. 18). Upon integrating spatial data, pathway enrichment analysis continued to highlight CellAgentChat’s superiority over CellPhoneDB and CellChat (Fig. 7c). CellAgentChat continues to perform better than CellPhoneDB in all datasets (Fig. 7c, Supplementary Fig. 18b). CellAgentChat also demonstrated superior performance to CellChat, NICHES COMMOT in the majority of datasets (Fig. 7c, Supplementary Fig. 18b). Overall, pathway enrichment analysis supported the CCI inference precision of CellAgentChat over other benchmarked methods.

Finally, we calculated the overlap in LR pairs among the top 100 identified LR pairs across the six methods. We observed that the overlap between CellAgentChat and the other methods, particularly CellChat, NICHES and COMMOT was high (Supplementary Fig. 19). This serves as another confirmation of our method’s performance, indicating that the LR signaling predictions made by CellAgentChat align with those of other state-of-the-art methods.

Building on the findings from the preceding benchmarking sections, which underscore CellAgentChat’s superior, or at least comparable performance compared to benchmarked methods, it is crucial to recognize that beyond these achievements, CellAgentChat also introduces distinct functionalities not available in other tools. Unique to CellAgentChat is its integrated ABM animation platform, which facilitates the visualization of individual-level cellular interactions and supports dynamic simulations. Additionally, through easy and flexible adjustment of agent behavior, CellAgentChat empowers the use of in silico receptor blocking to explore potential interventions effectively and serves as a valuable tool for identifying both long and short-range interactions.

## DISCUSSION

We introduced CellAgentChat, an agent-based model (ABM) capable of elucidating cell-cell interactions (CCI) from single-cell data, both with and without spatial information. As a novel approach, CellAgentChat employs individual cell agents guided by simple behavior rules to investigate the complexity of cellular interactions within the system. In contrast to the existing methods for predicting CCI, CellAgentChat offers three unique features. Firstly, it offers a dynamic visualization of cellular interactions related to individual cell agents. Through its cellular interaction animation, one can explore cellular heterogeneity within a population, often uncovering insights at the cluster level that are impossible to obtain (Fig. 5). Thus, CellAgentChat provides both a cluster- and cell-specific view of cellular communication. Secondly, CellAgentChat enables efficient in silico receptor blocking, facilitated by modulating the agent rules (for instance, receptor receiving rate), allowing for a systematic assessment of a receptor blocking’s impact on downstream gene expression dynamics. CellAgentChat can propose systematic strategies for potential therapeutic interventions by investigating disease-related genes.

Using CellAgentChat, we identified key LR pairs known to play crucial roles in neural development and cancer. We identify the *Apoe* and collagen signaling pathways among microglial and vascular cells in mouse neurogenesis (Fig. 2). Both pathways have been shown to play critical roles in neurogenesis^33–36^. Furthermore, we identified key time-dependent signaling pathways such as radial glial cells (RGCs) role in producing intermediate progenitor cells and neurons. CellAgentChat also revealed a significant *LAMC2-ITGB6* interaction among tumor keratinocytes (TKer cells) in squamous cell carcinoma (Fig. 3). This interaction has previously been linked to critical roles in tumor progression and migration, suggesting its involvement in the early stages of cancer development^55^. In addition, CellAgentChat confirmed the selection of three receptors, *EGFR*, *PDCD1* (*PD-1*) and *CTLA4*, among others for receptor-targeting treatment options in breast cancer (Fig. 4). Thirdly, researchers can tailor CellAgentChat to prioritize long- or short-range interactions controlled by the diffusion rate employed in quantifying cellular interactions. Using this approach, we identified *Sema4a-Plxna4* as a short-range interaction involved in axon-guidance in the mouse cortex^69^ (Fig. 6). Finally, we showed that CellAgentChat surpasses several state-of-the-art methods across multiple metrics, including the generation of cell-cell communication networks, alignment with spatial information, Reactome pathway enrichment analysis, and overlap of ligand-receptor pairs across different methods (Fig. 7).

In future work, we consider the following extensions of CellAgentChat. By representing cells as individual agents with distinct statuses, ABMs enable precise modeling of heterogeneous cell populations, effectively capturing the diversity in cell types, states, and behaviors essential for comprehending biological phenomena^27,28^. Expanding CellAgentChat’s capabilities to include cell movement and even cell death would be natural extensions, enhancing its capacity to simulate realistic scenarios. Although technically challenging, further improvements by incorporating ligands, receptors, and other cofactor molecules as agents in addition to cells would allow for a more comprehensive and detailed simulation of biological systems’ complex interactions and dynamics. By considering these additional components as agents, the model could capture the intricate interplay between cells and their surrounding molecular environment, providing a more nuanced understanding of the underlying mechanisms at play. It would also improve the efficiency of in silico perturbations such as receptor blocking. Given the prevalent issue of low gene coverage in many single-cell resolution spatial datasets, like those generated with SeqFISH+ and Xenium, spot-level technology remains a popular choice among researchers. In our framework, an agent can represent individual cells or spots, adapting for analytical flexibility. While this representation may overlook potential interactions between multiple cells within the same spot, especially since many CCIs occur within a 250µm range^71^, our approach simplifies the analysis of complex CCIs in spot-level data and demonstrates superior interaction inference compared to other methods. Users inclined towards analyzing spot-decomposed cell interactions, rather than staying at the spot level, can employ tools such as SpatialScope^72^ to deconvolve spot-level spatial transcriptomics data, achieving single-cell resolution. With this enhanced resolution, CellAgentChat can be run to explore and understand CCI dynamics in greater detail.

## METHODS

### Ligand-Receptor database

In this study, we used the ligand-receptor (LR) interactions from CellTalkDB^73^, a curated database of almost 3400 human and over 2000 mouse literature-supported interactions, as the default LR interaction resource for our CellAgentChat method. The interactions from the curated database specify a comprehensive collection of expert curated LR interactions. Besides the default CellTalkDB LR interactions, CellAgentChat also allows for customized LR interactions that users obtain from other sources or manually curate.

### Single-cell sequencing data preprocessing

We preprocessed the single-cell RNA-sequencing (scRNA-seq) data using the Scanpy^74^ pipeline. We began by filtering out cells with more than 20% mitochondrial reads, which are indicative of poor-quality cells^75,76^. We also removed genes not expressed by a minimum of 3 cells. Finally, cell counts are normalized against the library size and transformed into log space (with the *log1p* function). As an ablation study, we show that using normalized gene expression values provide better results than using raw unprocessed counts (Supplementary Fig. 20). The spatial coordinates of each cell are incorporated into the “anndata” object (in h5ad format) with the gene expression.

### Scoring cell-cell interactions with the agent-based model

To build the agent-based model (ABM), we need to define the states and behavior of each agent. Agents here represent the cells in the biosystem. The agent states describe the state and features associated with each cell, while the agent rules define how each agent behaves.

### Cell Agent State Description

As we are primarily interested in studying the interactions between cells (or clusters), we describe each cell agent with characteristics and features related to cellular interactions. The remainder of this section provides the detailed cell agent state description. (1) *Cell Agent Expression*: The expression value of all genes, including ligands and receptors, is stored for each cell. The gene expression profile presents the quantitative description of the cell agent state, which is essential to infer the interaction behaviors. (2) *Cell Agent Spatial Position*: We use spatial transcriptomics data to record the position of each cell agent. All the cells are placed on the grid at their spatial coordinate (if provided). Otherwise, CellAgentChat randomly distributes all cells on the grid. (3) *Cell Agent Batch*: To utilize multiple samples (biological replicates or technical samples) concurrently, we designate the batch of each cell to confine interactions solely between cells of the same batch. (4) *Cell Agent Ligand-Receptor Universe*: Every cell agent has the same LR universe using interactions curated from CellTalkDB or user-customized LR interactions. Users can modify the LR Universe by blocking a specific receptor or LR interaction (see Agent-Based Animation of CCIs). By doing so, they can remove all interactions associated with ligands or receptors (or specific LR interactions) from the universe of all cells. This unique characteristic of ABMs enables effective and flexible ligand/receptor perturbations.

### Cell Agent Communication Behavior Rules

In this study, the cellular communications between agents (cells) depend on the states of the cells, which includes the gene expression of the cell agent, the spatial position of the cell agent (optional) and the LR universe. To be more specific, the cellular interaction strengths (scores) between cell agents are quantified based on the set of defined rules as discussed below.

#### (1) Ligand Diffusion Rate 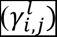

The diffusion rate 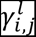 estimates the rate of the diffusion of a ligand *l* in the microenvironment from sender cell *i* to receiver cell *j*. The diffusion rate depends on a few factors. First, the diffusion rate is proportional to the number of ligands a sender cell secretes into the environment. We can estimate this from the number of RNA molecules proportional to its gene expression.

Second, the diffusion rate is inversely proportional to the distance (of the receiver agent) from the sender cell agent. We calculate the Euclidean distance between two cells based on the spatial coordinates of all the cell agents. The hyperparameter τ, is a dimensionality parameter representing the degrees of spatial freedom for spatial data. τ is typically set to two for spatial transcriptomics data derived from 2-dimensional slices. However, with the advancement of technology and the increasing prevalence of 3-dimensional slices, we can adjust the τ parameter accordingly (τ = 3)^77^. When spatial information is unavailable (e.g., in single-cell RNA-seq data), *τ* is set to 0 so that the distance between all cells in the data will degenerate to 1 (*d*^*τ*=0^ = 1).

In the absence of precise information about the rates of ligand travel or the operational distance range of LR pairs, we employ the delta (δ) parameter as an approximation for the distance range of LR pairs and model the decay rate of ligands. A user-adjustable slider δ (between *0-10*, default = *1*) is available to control the decay rate of the diffusion along with the increase of distance, which allows for inferring long-range or short-range interactions. Keeping δ at the default value of 1 provides no bias toward long-range or short-range interactions. We modeled short-range interactions with a δ = 10 and long-range interactions with δ = 0.1. By investigating various δ values and the corresponding agent communication behaviors defined by these values, we can also determine a specific δ for each LR pair. This approach avoids the assumption that all ligands share the same diffusion rate. The distance between sender cell *i* and receiver cell *j* is calculated as the Euclidean Distance *d*(*i*, *j*). The expression of ligand *l* in sender cell *i* is denoted as 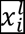. Then, the diffusion rate 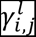 can be calculated as below:

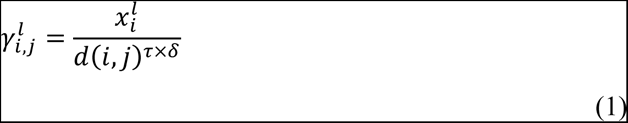

#### (2) Receptor receiving rate 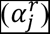

The receiving rate 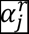 associated with a receptor *r* represents the probability that an interaction is received by the receptor in cell *j*, and it is proportional to the expression of the receptor. The receptor receiving rate will be a value in the range (0,1) where 0 represents the receptor cannot “receive” any ligands that reach it, while 1 denotes that reached ligands will all be “received” by the receptor. Here, we employed the non-parametric min-max normalization to calculate the receptor receiving rate for each receptor. Min-max normalization has been used in other methods for cell interaction inference^78–81^, lending support to our choice. For each cell, the expression value of each receptor is subtracted by the minimum expression value in that cell and then divided by the difference between the maximum and minimum expression values in that cell (min-max normalization), enabling the identification of highly expressed receptors within each cell, thereby establishing a cell-specific metric.

#### (3) Receptor transcriptome conversion rate (*β*^*r*^)

CCIs play a crucial role in the regulation of gene expression, impacting the transcriptome, and ultimately influencing cellular function and fate. In order to account for the effect of cellular interaction on downstream gene expression, we employed a neural network regressor to estimate the downstream gene expression impact associated with each receptor (here we term it as the receptor conversion rate *β*^*r*^). The neural network takes as input two vectors, 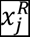, representing expression of all receptors in cell *j* and 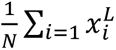, representing the average expression of all ligands across the set of all *N* cells. These two vectors are then fed into the neural network *(f)* to predict the expression 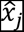 of all genes in cell *j* (regression task). The neural network structure comprises the following components: two input layers, two hidden layers, and one output layer. The size of the two input layers corresponds to the number of ligands and receptors, respectively. The first hidden layer is dedicated to representing specific LR pairs. Each node in this layer receives two connections, one corresponding to a ligand and the other to a receptor, thus encapsulating the unique LR pair interactions. The second hidden layer represents transcription factors involved in downstream signaling pathways. We leverage scSeqComm^26^, a database that incorporates insights from KEGG^82^ and REACTOME^83^ pathways to delineate the association between a receptor and its downstream transcription factors (TFs), to inform the connections between this layer and the previous one. The output layer is responsible for outputting downstream gene expression predictions. To establish connections between known TFs and their downstream gene targets, we utilize prior knowledge from databases such as TRRUST v2^84^, HTRIdb^85^, and RegNetwork. If specific information regarding the target genes of certain TFs is unavailable, we utilize dense connections. We use a stochastic gradient descent optimizer to train the network ℱ that minimizes the mean squared error loss *L* (between the true observed expression for the set of all cells *X* and the predicted expression):

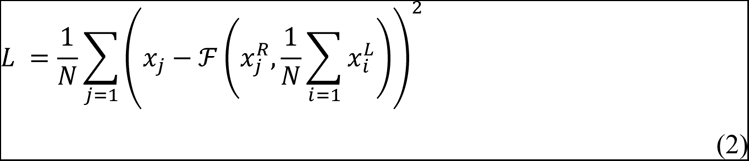

With the trained neural network regressor, we then perform a sensitivity analysis to score each input receptor based on its impact on the downstream gene expression. Specifically, we conduct a perturbation-based feature calculation^86^, by permuting feature values for each receptor, independent of the output. Next, we adjust the expression of the permutated receptor to reflect a ∼50% reduction, scaling it by the median expression of all receptors. This represents the perturbed input receptor vector (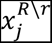) simulating a ∼50% repression of receptor *r* in expression. While this 50% repression serves as the default setting, we also offer the option to simulate a 100% repression, equivalent to a complete knockout, catering to users who prefer a full knockout scenario. Then, we quantify the importance of the receptor *r* (*I*_*r*_) based on the loss increase caused by the perturbation:

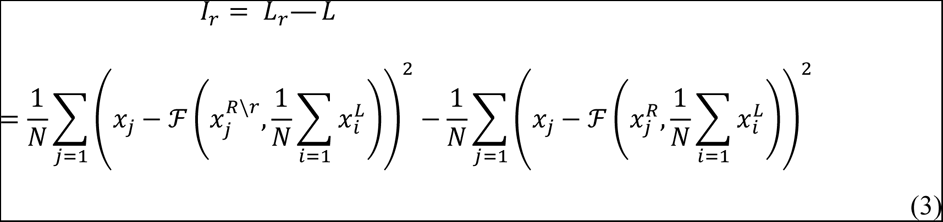

Employing a similar approach as the receiving rate, we normalize the receptor importance into range (0,1) via min-max normalization. Here, we set a minimum value for the conversion rate to be 0.4 based on previous studies that measure the effect of cellular interaction on gene expression^87^. We confirmed that this approach does not significantly alter the resulting gene expression of the cells of the training grid (Supplementary Fig. 21 and 22).

#### (4) Cell-cell interaction score between cells

The interaction score between a ligand-receptor pair (*l*, *r*) in a sender-receiver pair (*i*, *j*) is calculated based on the rates calculated above (*γ*^*l*^, *α*^*r*^, *β*^*r*^):

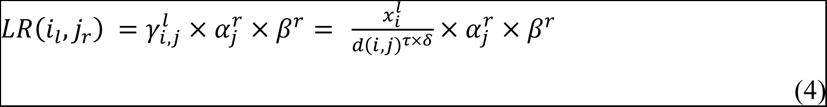

The cell receiving score for a given cell *i* is calculated as the summation over all ligand-receptor pairs (l,r) across all sending cells:

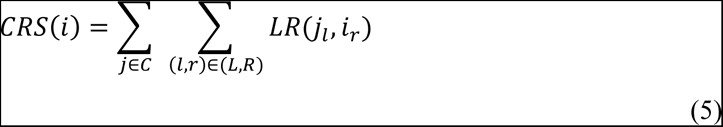

The combined interaction score between a sender-receiver cell pair (*i*, *j*) is calculated as the summation over all ligand-receptor pairs (l,r):

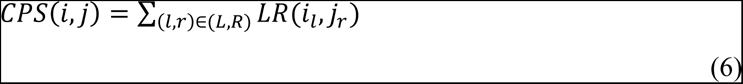

The overall interaction score *IS*(C1, C2) between two clusters C1 (sender) and C2 (receiver) for a ligand-receptor pair (l,r) is quantified as the average interaction score of all possible sender- receptor cell agent pairs between the sender C1 and receiver C2:

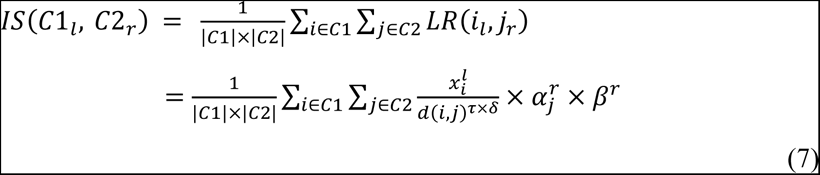

We also introduce a method to optionally integrate the utilization of pseudotime ordering of cells into the calculation of CCI scores. Please refer to the supplementary text for further details.

### Finding significant ligand-receptor pairs between cell populations

Once we get the interaction score for an LR pair between two cell clusters, we will examine whether such an interaction score is statistically significant (stronger compared to the random background). Specifically, we employed a random permutation test to calculate the statistical significance P-value associated with each LR interaction score. Significant interactions are identified between two cell groups using a permutation test. It randomly permutes the group labels of cells and then recalculates the interaction score for each LR pair. The LR score is stored after each permutation regardless of the cell type pairing. We perform enough permutations so that each LR pair, has on average 10,000 score values. We then obtained the maximum likelihood estimates of alpha and scale of Gamma distribution from the random scores for each LR pair. Using the background distribution, we can compute a P-value for a given interaction score (for the specific LR pair) by evaluating its position in the distribution’s right tail. The P-value is then corrected using FDR (Benjamini/Hochberg) correction. Interaction Scores (IS) with a P-value above the threshold (0.05) are considered Significant Interaction Scores (SIS). Using the SIS, we can calculate the cell type pair scores (CTPS) between two clusters, C1 (sender) and C2 (receiver):

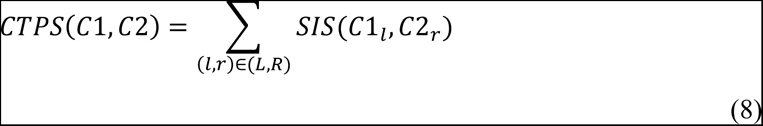

Due to the computational expense of calculating the distances between all cells in the dataset, the permutation test can be quite costly when considering spatial information. To address this, we have devised a method to match the background gamma distribution of non-spatial calculations with that of spatial ones. This is achieved by dividing the scale parameter by the average distance between all cells. Additionally, we halve both the alpha and the modified scale values. Although this does not equate to perfect scaling, we present very similar results using the scaling method significantly more efficiently (Supplementary Fig. 23b, c).

### In silico receptor perturbation

After identifying the top interactions between cell types, we conducted in silico perturbation using our trained deep learning model *f*. To illustrate the process of in silico receptor blocking, we centered our attention on the receptors involved in the top 25 LR interactions, spanning all cell type pairs. We then subjected these receptors to a perturbation analysis.

We permutated feature values for each receptor, independent of the output and simulated receptor blocking and scaled the expression of the permutated receptor by the median expression of all receptors. This manipulation of the input receptor vector, 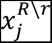, allowed us to gauge the ensuing influence on the gene expression downstream. After each receptor’s perturbation, we singled out the most impacted genes on average across all cells. Specifically, we looked at the top 50 most impacted genes. This approach enabled us to assess the effects of receptor blocking at the transcriptome level. To further demonstrate the potency of our model in executing in silico perturbation, we utilized the DisGeNET^57^ database as the potential validation. We verified whether the perturbed genes are associated with the specific diseases that pique our interest – Breast Cancer/PDAC (i.e., the disease associated with the dataset). To quantify the strength of the association between each receptor and the disease, we applied a binomial test to compute the statistical significance of the overlap between the perturbed genes and the genes associated with each disease (breast cancer, breast carcinoma, PDAC, pancreatic carcinoma). We further refined our results by adjusting the P-values over all disease gene sets, utilizing the FDR-Bonferroni Correction method, ensuring the robustness of our findings. Additionally, we confirmed the effectiveness of these receptor-inhibitor therapeutics using Kaplan-Meier (KM) survival plots^88^.

### Agent Based Dynamic Simulations and Animation of CCIs

CellAgentChat offers dynamic simulation capabilities for analyzing the impact of CCIs on cell states over short durations. Users have the flexibility to either execute the model once, emulating other state-of-the-art methods and inferring CCIs at the current moment, or they can run it over multiple short time steps to simulate temporal cell state dynamics. At each time point, we compute CCIs as described earlier. Subsequently, our trained neural network predicts gene expression levels for each cell post-CCI for the next time step. Although we cannot directly substitute existing gene expression values for the next time step with predicted ones due to uncertainties since many factors other than CCIs could have an influence on the expression, we place trust in the directional changes for each gene —whether they increase or decrease^70^. For each cell we then identify genes with statistically significant changes and adjust their expression for that cell within the ABM, either increasing or decreasing them by a small, specified amount (0.01) to simulate minor expression changes^70^. This iterative process is repeated across all time steps, reflecting the dynamic modulation of gene expression by CCIs.

CellAgentChat offers a unique feature by allowing the inference of cellular interactions between individual cells rather than just cell types. Its real-time animation platform displays the CRS of each cell, with size representing the strength; larger cells indicate stronger CRS (Supplementary Fig. 23a). The platform also includes customizable hyperparameters, such as the maximum number of steps (default of 1), δ for controlling long-range and short-range interactions (default of 1) and the degrees of spatial freedom τ (default of 2) enabling effective in silico permutations. See supplementary text for more information on how to effectively set up the hyperparameters to facilitate simulations.

In addition, CellAgentChat offers hyperparameters for controlling the LR universe. Users can block a specific LR pair or a specific receptor to observe downstream gene expression changes, which can aid in identifying potential therapeutic targets. The animation platform also allows for visualizing specific cell types/clusters, with modifications only applied to those selected. Furthermore, users have the option to visualize specific receptor or LR pairs. This visualization tool enables researchers to understand cell communication patterns at a granular level.

### Functional Evaluation of Ligand-Receptor pairs inferred by CellAgentChat

We conducted pathway enrichment analysis to validate the interactions inferred by CellAgentChat functionally. We utilized both ligands and receptors from the top 25 highest scoring LR pairs among all cell type pairs as the input gene set. To perform pathway enrichment analysis, we employed g: Profiler^89^ and selected all terms from the Reactome pathway with a minimum adjusted P-value below 0.01. This approach was consistent across all datasets, regardless of spatial or non-spatial. In the cases where we compared the enrichments for the Reactome pathways between two groups (e.g. long-range group vs. short-range group), we used signed-rank Wilcoxon test.

### Benchmarking

We conducted a comparative analysis of CellAgentChat with five other state-of-the-art methods, namely CellChat, CellPhoneDB, NICHES, COMMOT, and Scriabin. Among these, CellChat, CellPhoneDB, and NICHES all facilitate the inference of cell-cell interactions (CCIs) using both spatial and non-spatial data. Conversely, COMMOT exclusively supports spatial data, while Scriabin exclusively supports non-spatial data. Additionally, Scriabin allows for the computation of a cell-cell interaction matrix (CCIM); however, it is recommended only for small datasets due to scalability limitations. Consequently, we could only evaluate Scriabin’s performance for the PDAC and the SeqFISH+ mouse somatosensory cortex datasets. We used the LR universe derived from CellTalkDB for all methods.

We conducted systematic benchmarking with and without spatial data to validate CellAgentChat’s ability to accurately infer interactions prior to integrating the spatial tendencies of CCIs. To assess the accuracies of the methods, we measured the Pearson Correlation to evaluate the communication pattern across cell types of each of the six methods. In the absence of a ground-truth, we employed an ensemble approach. We normalized the interaction matrix computed by each method and took the average to create an ensemble interaction matrix as our new “ground-truth”. We then calculated the Pearson correlation between the ensemble interaction matrix and the inferred communication patterns of each method. Additionally, we compared the interaction matrix of each method with the overall distance matrix of the cells to verify whether CellAgentChat’s CCI predictions were influenced by the spatial proximity between cells. Specifically, we calculated the Pearson Correlation between the cell interaction matrix with the cell-cell inverse distance (1/*d*) matrix.

As described above, we also conducted pathway enrichment analysis using the genes from the top 100 LR pairs to evaluate the functional validation of the LR interactions inferred by all six methods. We identified pathways through Reactome pathway enrichment analysis of the inferred interactions by at least two methods, to capture the most relevant pathways, and compared the - log_10_ adjusted P-values across all methods. To differentiate the functional validity of the LR interactions across the methods we used the Wilcoxon test (one-sided).

Finally, to compare the similarity in the inferred interactions we determined the number of shared LR interactions from the top 100 pairs identified by each method.

## Data availability

CellTalkDB is available at (https://github.com/ZJUFanLab/CellTalkDB). The SCC and PDAC datasets analyzed in this study are available from the Gene Expression Omnibus (GEO) repository under the following accession numbers: GSE144240 (GSM4565826) and GSE111672 (GSM3036911), respectively. The data for the SeqFISH+ mouse somatosensory cortex dataset can be found at the following GitHub link: https://github.com/CaiGroup/seqFISH-PLUS. The Xenium human breast cancer dataset (sample 2) can be found at https://www.10xgenomics.com/products/xenium-in-situ/preview-dataset-human-breast. The stereo-seq developmental mouse hippocampus dataset can be found at STOmicsDB^90^ https://db.cngb.org/stomics/mosta/. We used the h5ad file titled “*Dorsal_midbrain_cell_bin.h5ad”*.

## Code availability

The CellAgentChat software package and source code are available at our GitHub repository (https://github.com/mcgilldinglab/CellAgentChat). All preprocessed .h5ad files used in this study are also available in the same GitHub repository.

## Acknowledgements

This work is supported by grants from the Canadian Institutes of Health Research (CIHR) [PJT- 180505 to J.D]; the Fonds de recherche du Québec - Santé (FRQS) [295298 to J.D., 295299 to J.D.]; the Natural Sciences and Engineering Research Council of Canada (NSERC) [RGPIN2022- 04399 to J.D.; RGPIN-2016-05174 to Y.L.]; and the Meakins-Christie Chair in Respiratory Research [to J.D.]. This publication is part of the Human Cell Atlas (HCA) publication bundle (HCA-10).

## Notes

### Competing Interest Statement

The authors have declared no competing interest.

### Summary of Updates

Improved modeling parameters. Addition of several spatial transcriptomics datasets at single-cell resolution. More comprehensive benchmarking of CellAgentChat with other state-of-the-art methods.

