## Supplemental Text for "Harnessing Agent-Based Modeling in CellAgentChat to Unravel Cell-Cell Interactions from Single-Cell Data"

### SUPPLEMENTARY INFORMATION

#### Supplementary Text

##### CellAgentChat in silico receptor blocking for analysis of perturbed downstream genes in PDAC

To supplement the use of in-silico perturbation results on human breast cancer data at high spatial resolution, we also used CellAgentChat to conduct in-silico perturbation on a spot level PDAC dataset. Following the same procedure as for the breast cancer analysis, we employed CellAgentChat to identify key receptors involved in LR interactions and simulated the blocking of these receptors (Supplementary Fig. 4e, f). This led to the identification of the top 50 perturbed genes most impacted by each receptor's blocking. The DisGeNET enrichment analysis to evaluate the potential therapeutic benefits of receptor blocking identified four candidate receptors, *NOTCH3*, *PLAUR*, *ITGA11* and *CD74*, which exhibited the highest -log10 P-values (binomial test, FDR corrected, <0.05) for PDAC (Supplementary Fig. 8a). Additional analysis revealed that, when blocked, *CD63*, *NOTCH3*, *ERBB2*, *PLAUR*, *ITGA11* and *CD74* yielded statistically significant P-values (binomial test, FDR corrected, <0.05) for a disease similar to PDAC, Pancreatic Carcinoma (PC) (Supplementary Fig. 8a). The results without using the TF prior are shown in Supplementary Fig. 9b.

The cell receiving score (CRS) of each cell, as identified from our animation platform, depicts that Acin3, Strom, Duct, Can1, and Can2 cells exhibit the highest levels of involvement in interactions with the candidate receptors (Supplementary Fig. 8b). We have identified notable interactions among these populations, bolstering this conclusion (Supplementary Fig. 4e, f). While ductal cells traditionally serve as the primary origin of PDAC, emerging evidence suggests that acinar cells may also contribute to the development of PDAC[1]. This intriguing possibility could explain the extensive involvement of both Duct and Acin3 cells in interactions with all three receptors (Supplementary Fig. 8b). Furthermore, studies have demonstrated that stromal cells significantly influence extracellular matrix (ECM) formation and tumor progression in PDAC[2]. Through an agent-based view, we gain robust confirmation of the extensive involvement of Strom, Duct, Can1, and Can2 cells in the context of PDAC. Notably, the agent-based approach reveals even more pronounced interactions with Acin3 cells, further enriching our understanding of the intricate dynamics at play in this disease.

While studies have linked many of the receptors analyzed in this study with PDAC, the candidate receptors we identified — *NOTCH3*, *PLAUR*, *ITGA11* and *CD74*—appear to play a particularly crucial role in PDAC development[3-6] (Supplementary Fig. 8, 9). This finding validates the efficacy of the in-silico receptor blocking technique facilitated by CellAgentChat. Blocking the *NOTCH3*, *PLAUR*, *ITGA11* and *CD74* receptors in silico perturbs the disease genes associated with PDAC (from DisGeNET) (Supplementary Fig. 8c, e, g, i, 9c). In particular, when *PLAUR* was blocked, *DHX58* emerged as the fourth most affected gene, previously linked with PDAC[7] (Supplementary Fig. 9c). All other PDAC disease target genes perturbed by blocking the candidate receptors are depicted in Supplementary Figure. 8 and 9, highlighted in red.

**Integration of cell agent pseudotime in calculating cell-cell interaction score**

<u>Cell Agent State: Pseudotime</u>

Even the cells within the same cluster exhibit differences, in many continuous biological processes, such as cell differentiation or disease progression. Pseudotime provides ordering of the cells within the cluster (and the entire dataset). The interactions of cells with varying pseudotime may differ substantially, even if they belong to the same cluster[8]. Therefore, here we include the pseudotime as an cell agent state description. In order to calculate the pseudotime trajectory of each cell, we use Slingshot[9], a cell lineage and pseudotime inference tool. Using the pseudotime values generated from Slingshot, we group each cluster's cells into different pseudotime bins (“stages”). Following the framework developed by TraSig[8], CellAgentChat stores the pseudotime bin that each cell resides in and uses it to calculate the interaction score for the single-cell data with spatial information.

<u>Cell Agent Communication Behavior Rules with Pseudotime</u>

The CCI score discussed in the main text ignores the pseudotime time differences between all cell agents in the same cluster. However, the integration of the cell pseudotime improves the CCI inference[8]. The main difference in these score calculations lies in the fact that, instead of computing scores between every cell in two clusters, we now exclusively calculate scores between cells that reside within the same pseudotime bin ($b$):

$$CRS_{PT}(i) = \begin{cases} \sum_{j \in C} \sum_{(l,r) \in (L,R)} LR(i_l, j_r), & if\ b_i = b_j \\ 0 & , else \end{cases}, \quad (1)$$

$$CPS_{PT}(i,j) = \begin{cases} \sum_{(l,r) \in (L,R)} LR(i_l, j_r), & if\ b_i = b_j \\ 0 & , else \end{cases}, \quad (2)$$

We take the average score of all non-zero bins to produce a final pseudotime-integrated interaction score ($IS_{PT}$) for a given ligand-receptor (l,r) pair between two clusters:

$$IS_{PT}(C1_l, C2_r) = \begin{cases} \frac{1}{|C1| \times |C2|} \sum_{i \in C1} \sum_{j \in C2} LR(i_l, j_r), & if\ b_i = b_j \\ 0 & , else \end{cases}, \quad (3)$$

We also provide users with the choice to relax the constraint on cells interacting only when they are within the same pseudotime bin. We present a sliding window parameter (*w*) that allows for cells in neighboring pseudotime points/bins to interact. For example, if *w*=2, then a cell *i* in $b_i$ will interact with cell *j* in $b_j$ if $b_i = b_j$ or if $b_j$ is one of the neighboring two pseudotime bins in either direction from $b_i$. By default, *w*=0.

**Selection of optimal parameters for CCI inference and dynamic simulations with CellAgentChat**

Implementing CellAgentChat requires users to select several parameters. Here we discuss these parameters, their necessity in CCI calculation, their impact on obtained results, and suggestions for adjusting them to facilitate effective CCI inference and ABM simulations.

User defined hyperparameters essential for CCI inference with CellAgentChat include:

1) Tau ($\tau$): a parameter of the ligand diffusion rate that represents the degrees of spatial freedom concerning spatial distance (default, $\tau = 2$).
2) Delta ($\delta$): a parameter of the ligand diffusion rate that controls the decay rate of ligand diffusion (default, $\delta = 1$).
3) Sliding window (w): a parameter to control the restriction of cells in differing pseudotime (optional) bins from interacting (default, w = 1).
4) Time steps: The number of time steps/ticks the model will simulate (default = 1).

$\tau$ represents the degrees of spatial freedom concerning spatial data. $\tau$ is typically set to 2 for spatial transcriptomics data derived from 2-dimensional slices. However, as technology advances and 3-dimensional slices become more prevalent[10], we can adjust the $\tau$ parameter accordingly ($\tau$=3). When no spatial data is provided, $\tau$ is automatically set to 0 by the model.

$\delta$ controls the decay rate of ligand diffusion. Due to the lack of specific information regarding the rates at which each ligand travels or whether ligand-receptor pairs operate at short or long ranges, we set $\delta = 1$ for all ligands. However, we use the $\delta$ parameter as an estimation tool to infer the distance range of LR pairs. Our model can infer which LR pairs are long- or short-range. To effectively model these interactions, we recommend setting δ=10 for all ligands to simulate short-range interactions, and δ=0.1 for all ligands to simulate long-range interactions. This will allow the model to capture LR pairs in the data that work predominantly in long/short ranges. Using these results, we can then setup the ligand diffusion rate according to prescribe unique diffusion rates ($\delta$) for each ligand.

The sliding window parameter (w) governs whether cells in neighboring pseudotime points or bins can interact. By default, w=1, adhering to the methodology outlined in TraSig, which restricts cell interactions to those within the same bin. However, to address potential limitations arising from the assumption that cells with differing pseudotime values do not interact, we introduce the sliding window feature. This feature allows users to define a window within which interactions across pseudotime groups are permitted to a certain extent. We recommend using small values for w, such as w=1, to adhere closely to the TraSig approach or w=2 or w=3, to permit interactions across neighboring time points while still maintaining a degree of restriction to cells at similar time points.

The number of time steps the model can simulate is also customizable by the user. By default, the model runs for one time step, producing results reflecting the current cell states, similar to other

CCI inference methods. However, running CellAgentChat over multiple time steps enables users to analyze how CCIs modify cell states over time and influence subsequent interactions. This capability offers a deeper understanding of the dynamic nature of CCIs and their impact on cellular behaviors over time.

### SUPPLEMENTARY FIGURES

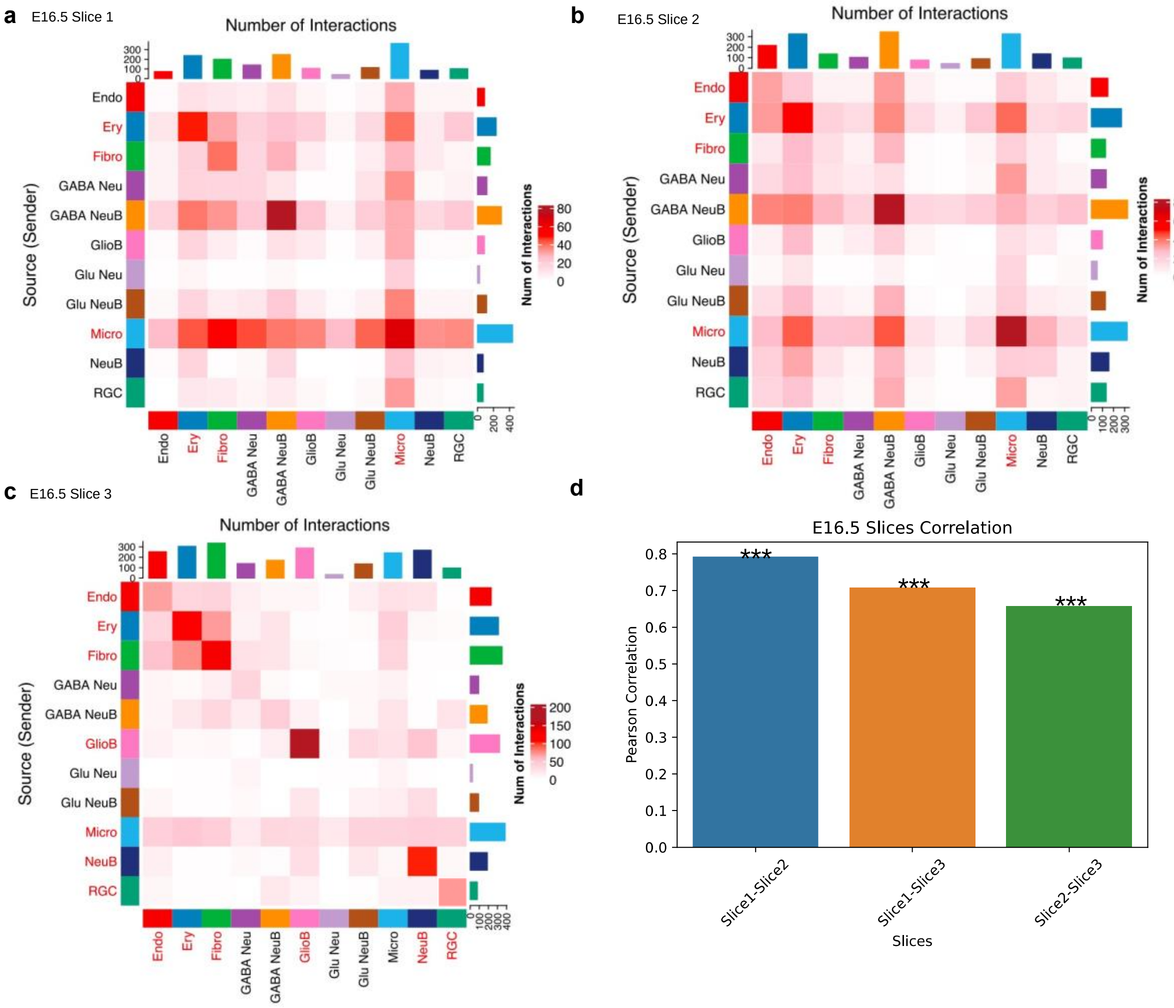


**Supplementary Figure 1: CellAgentChat can capture similar interactions across different technical samples.** (**a, b, c,**) Heatmaps display the communication network inferred by CellAgentChat in the developing mouse hippocampus at E16.5 across three different 2D spatial slices. **d,** Pairwise Pearson Correlation comparison of the communication pattern between each of the three slices.

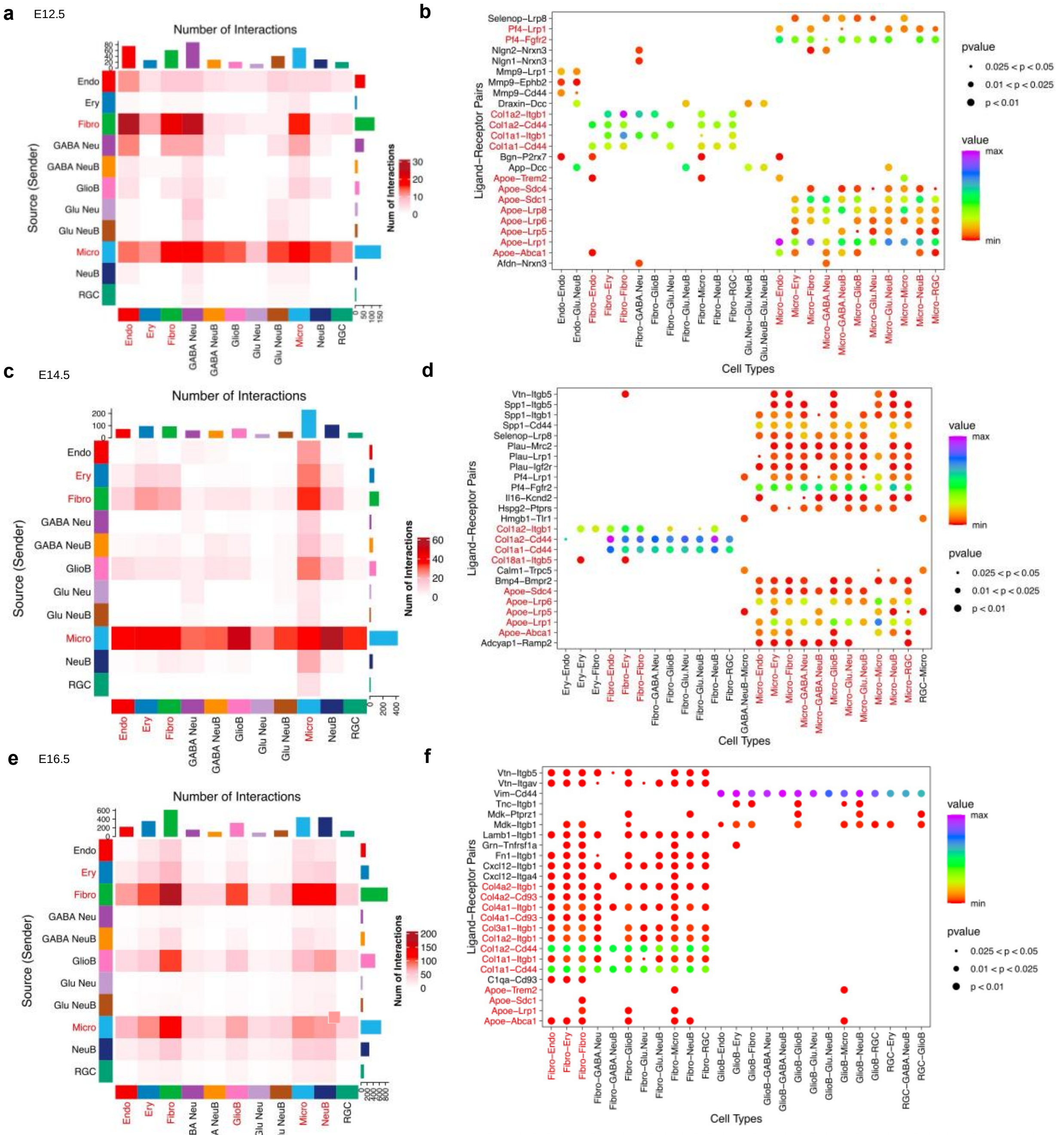


**Supplementary Figure 2: Analysis of Cell Communications in developing mouse hippocampus using non-spatial data.** (**a, c, e,)** Heatmaps display the communication network inferred by CellAgentChat at E12.5, E14.5 and E16.5, respectively. (**b, d, f,)** Dot plots illustrate the top 25 LR pairs between cell types inferred by CellAgentChat at E12.5, E14.5 and E16.5, respectively.

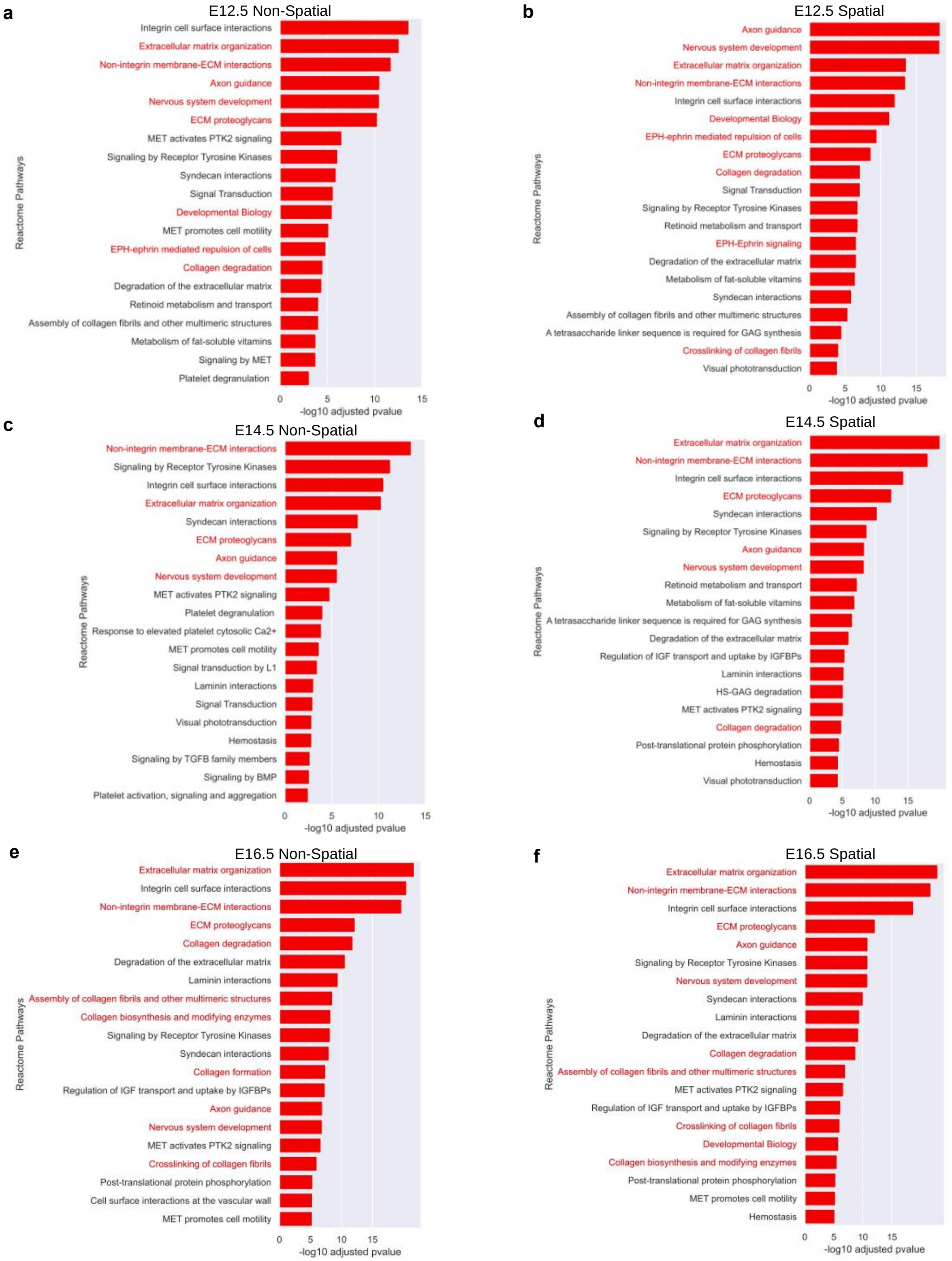


**Supplementary Figure 3:** Reactome pathways analysis using the ligand-receptor interactions inferred by CellAgentChat. **a,** Pathway analysis of E12.5 non spatial results. **b,** Pathway analysis of E12.5 spatial results. **c,** Pathway analysis of E14.5 non spatial results. **d,** Pathway analysis of E14.5 spatial results. **e,** Pathway analysis of E16.5 non spatial results. **f,** Pathway analysis of E16.5 spatial results. Red highlights indicate biological pathways relevant to neurogenesis.

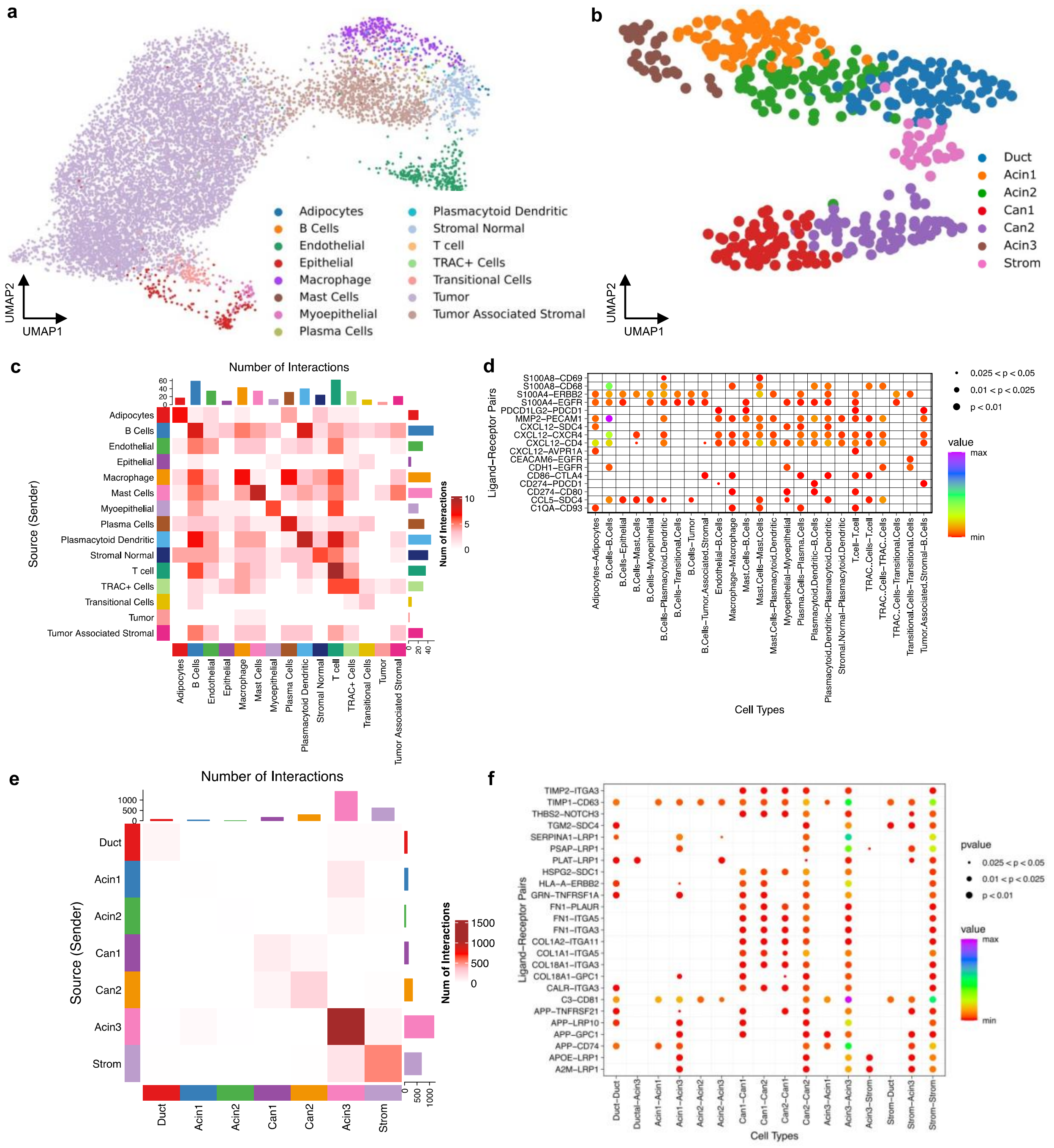


**Supplementary Figure 4: Inferred interactions by CellAgentChat on Human breast cancer and pancreatic ductal adenocarcinoma datasets a,** UMAP visualization of scRNA-seq data of Xenium human breast cancer dataset. **b,** UMAP visualization of scRNA-seq data of PDAC dataset. (**c, d,**) Heatmaps display the communication network inferred by CellAgentChat in breast cancer and PDAC, respectively. (**e, f,**) Dot plots illustrate the top 25 LR pairs between cell types inferred by CellAgentChat in breast cancer and PDAC, respectively.

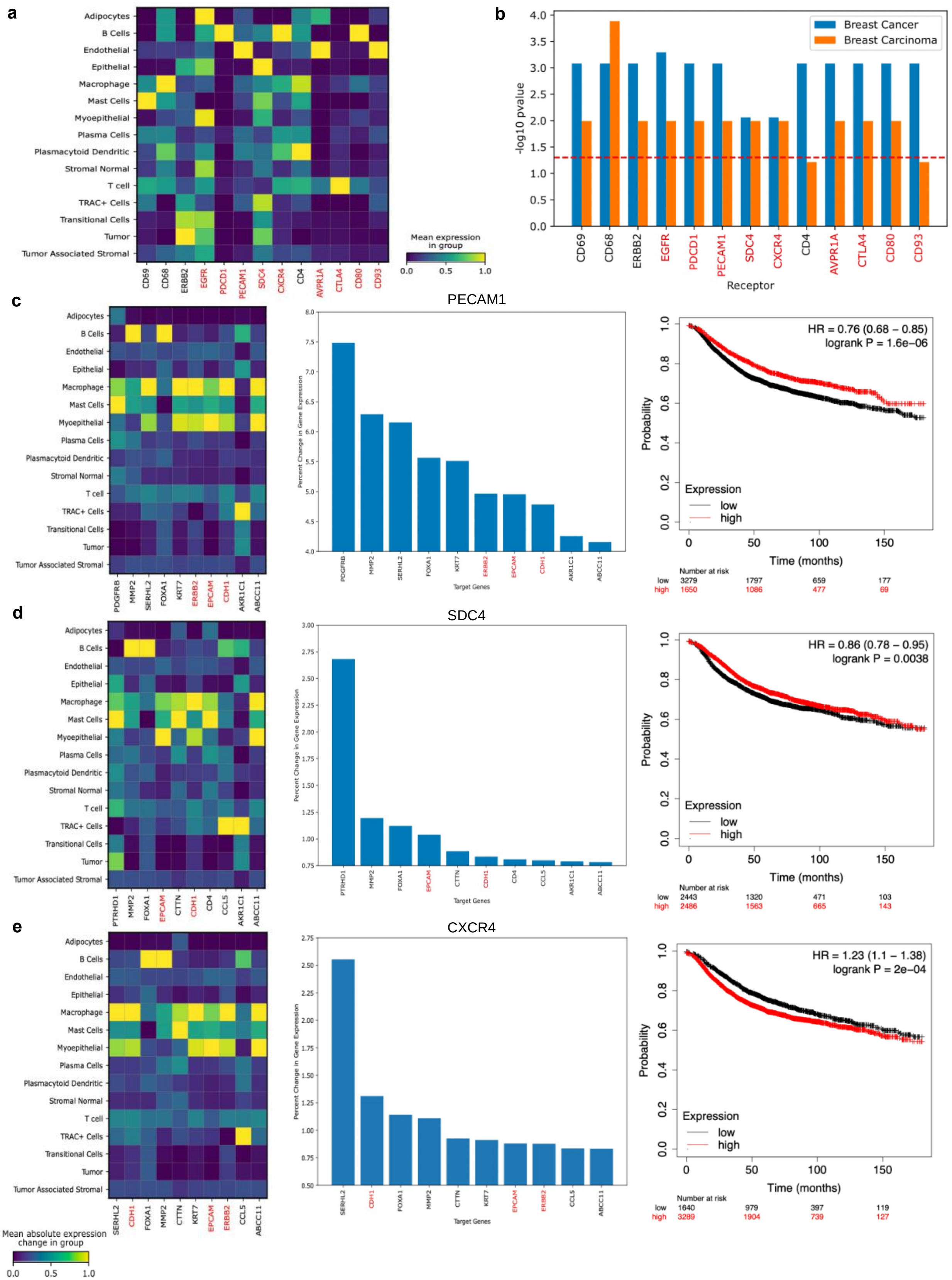


**Supplementary Figure 5: Identification of key downstream genes related to breast cancer after in silico blockage of top receptors. a,** Matrix plot shows the expression across cell types of the receptors involved in the top

25 ligand-receptor pairs inferred by CellAgentChat. **b,** Significance of receptor association with breast cancer and its similarity to a related disease (breast carcinoma), as represented by $-\log_{10}$ adjusted P-values (FDR corrected-Bonferroni Correction). Results are using the dense MLP without incorporating TF-prior LR-TF, TF-gene network information. (**c, d, e,)** Matrix plots display the mean change in expression of each target gene for each cell type for, *PECAM1*, *SDC4* and *CXCR4*, respectively (left). Identification of the top 10 downstream genes (target genes) with the highest percent change in absolute expression upon blocking *PECAM1*, *SDC4* and *CXCR4*, respectively (middle). A red highlight indicates genes associated with breast cancer. Survival analysis displaying the probability of survival of high vs. low receptor expression groups for *PECAM1*, *SDC4* and *CXCR4*, respectively (right).

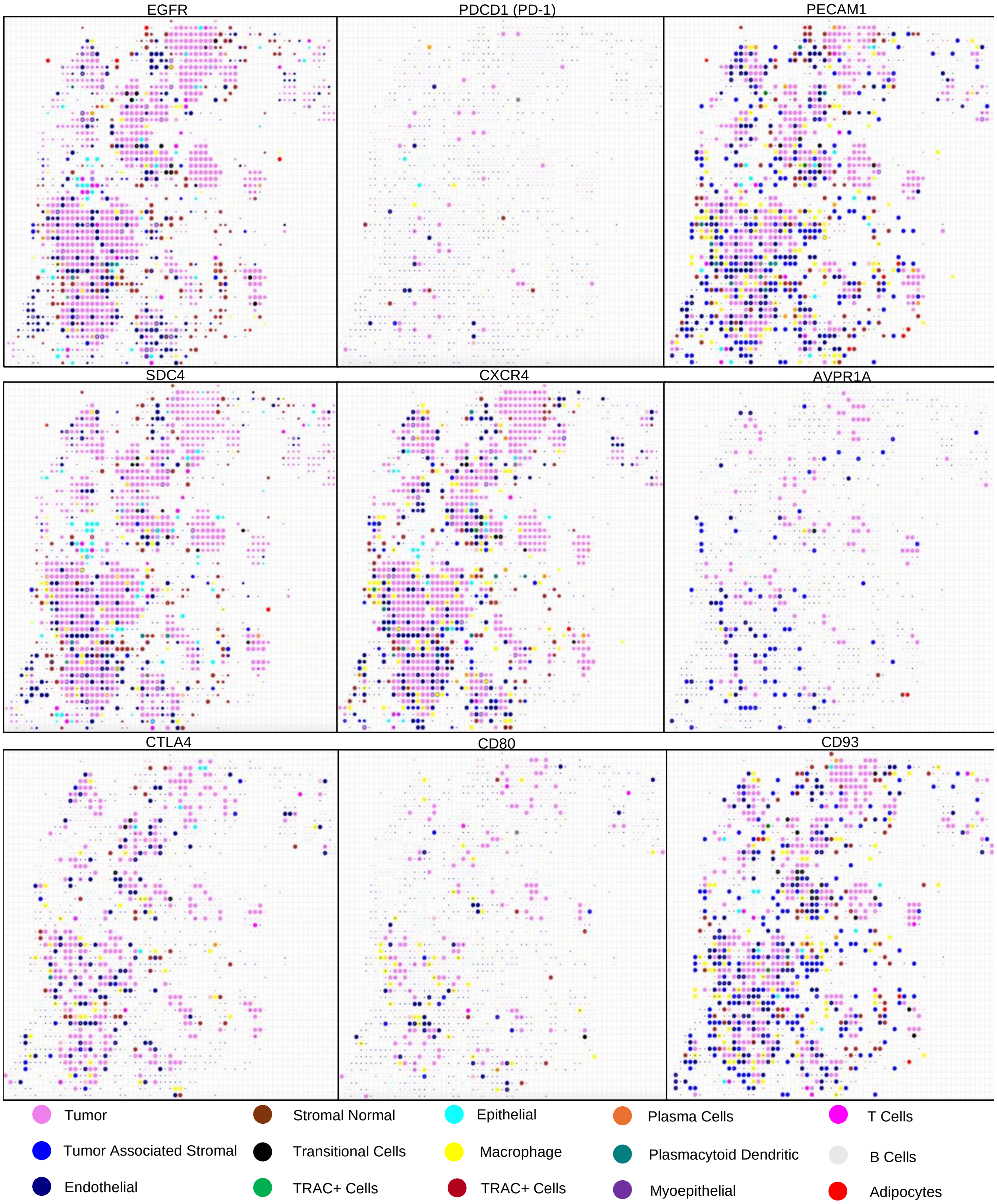


**Supplementary Figure 6: CellAgentChat's Animation depicts the cell receiving score (CRS) across individual cells for each candidate receptor.** After receptor blocking, all interactions involving these cell types will be completely blocked (go to zero), causing a shift in the interaction profile of the cells.

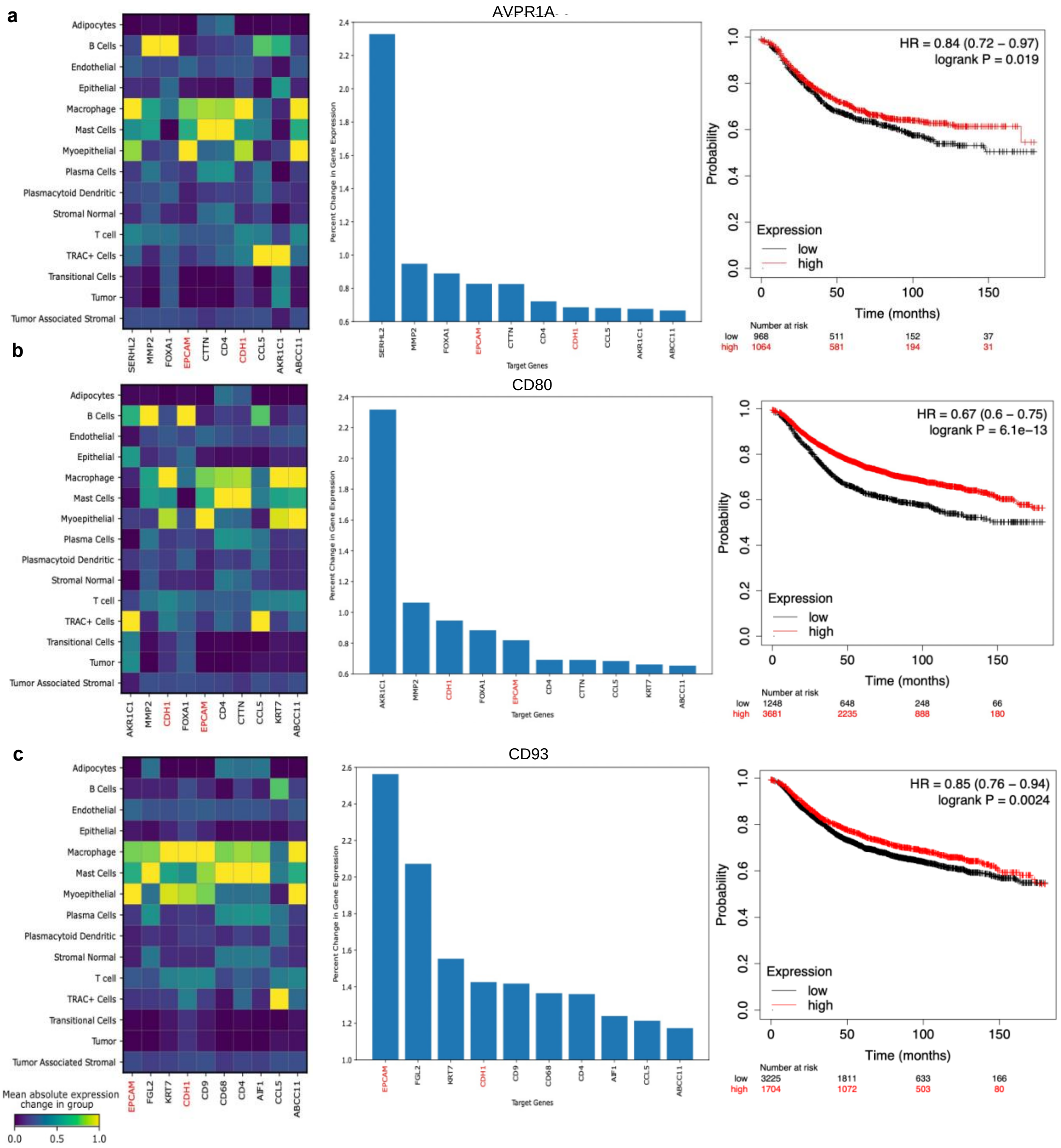


**Supplementary Figure 7: Identification of key downstream genes related to breast cancer after in silico blockage of top receptors.** (**a, b, c,)** Matrix plots display the mean change in expression of each target gene for each cell type for, *AVPR1A*, *CD80* and *CD93*, respectively (left). Identification of the top 10 downstream genes (target genes) with the highest percent change in absolute expression upon blocking *AVPR1A*, *CD80* and *CD93*, respectively (middle). A red highlight indicates genes associated with breast cancer. Survival analysis displaying the probability of survival of high vs. low receptor expression groups for *AVPR1A*, *CD80* and *CD93*, respectively (right).

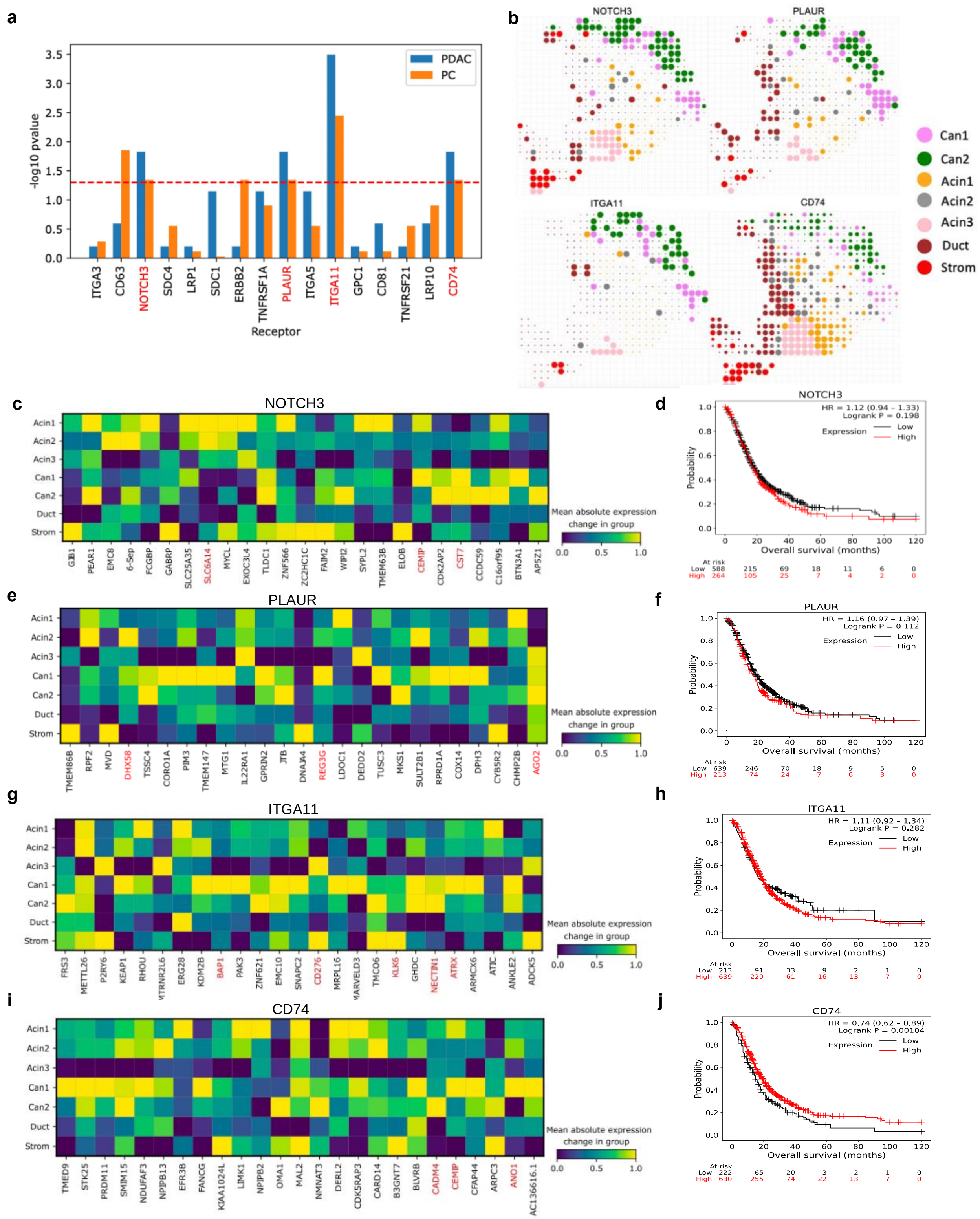


**Supplementary Figure 8: Identification of key downstream genes related to PDAC after in silico blockage of top receptors. a,** Significance of receptor association with pancreatic ductal adenocarcinoma (PDAC) and its similarity to a related disease, Pancreatic Carcinoma (PC), as represented by -$\log_{10}$ adjusted P-values (binomial test; FDR corrected-Bonferroni Correction). *NOTCH3*, *PLAUR*, *ITGA11* and *CD74*, highlighted in red, have significant -$\log_{10}$ adjusted P-values. **b,** The cell receiving score (CRS) for all interactions involving the receptors *NOTCH3*,

*PLAUR*, *ITGA11* and *CD74* determined using CellAgentChat's animation platform. The size of the cell in the animation indicates the CRS. **(c, e, g, i,)** Matrix plots display the mean change in expression of each target gene for each cell type for *NOTCH3*, *PLAUR*, *ITGA11* and *CD74*, respectively. A red highlight indicates downstream genes associated with PDAC. **(d, f, h, j,)** Survival analysis displaying the probability of survival of high vs. low receptor expression groups for *NOTCH3*, *PLAUR*, *ITGA11* and *CD74*, respectively.

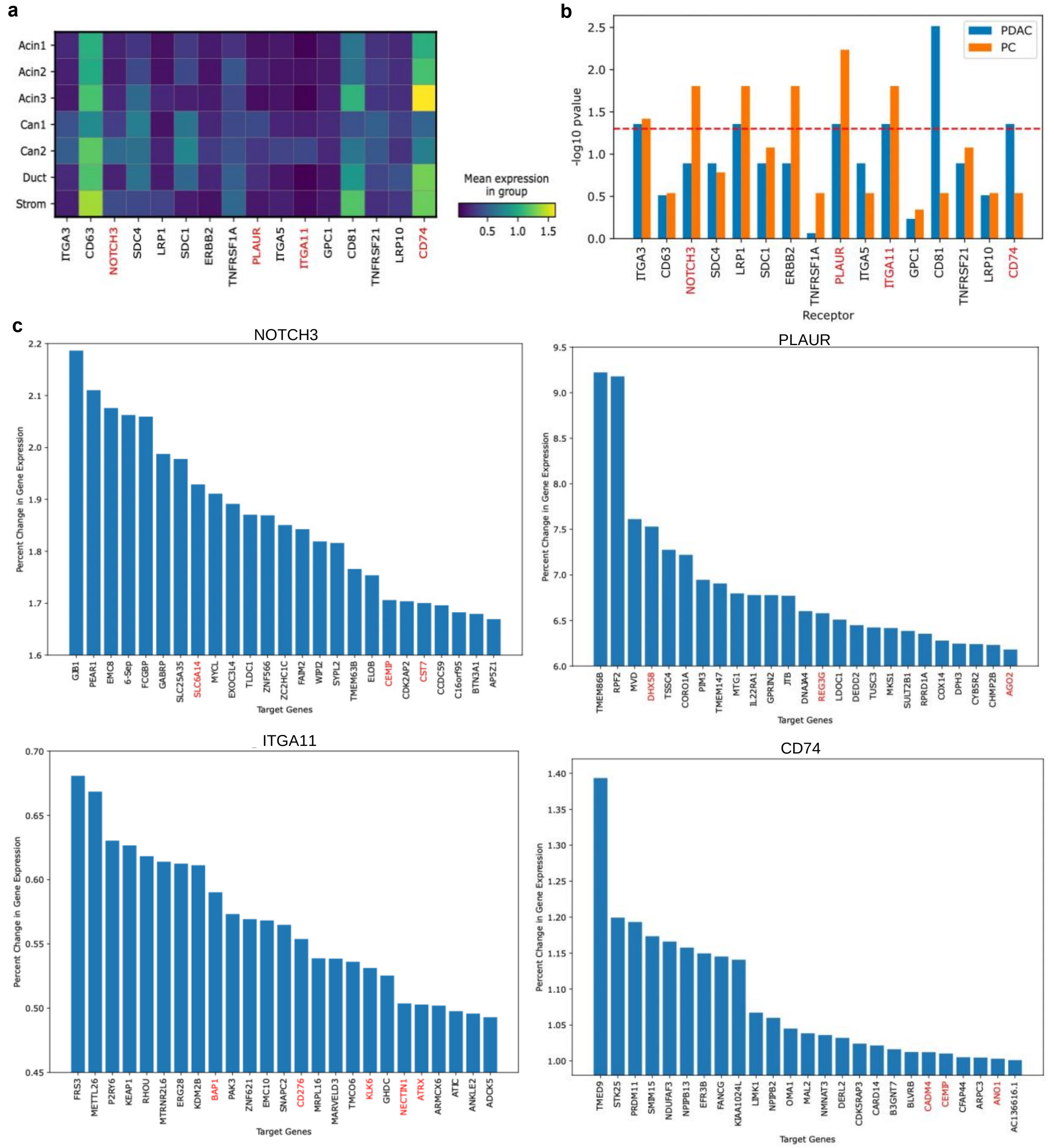


**Supplementary Figure 9: Identification of key downstream genes related to PDAC after in silico blockage of top receptors. a,** Matrix plot shows the expression across cell types of the receptors involved in the top 25 ligand-receptor pairs inferred by CellAgentChat. **b,** Significance of receptor association with PDAC and its similarity to a related disease (pancreatic carcinoma), as represented by -$\log_{10}$ adjusted P-values (FDR corrected-Bonferroni Correction). Results are using the dense MLP without incorporating TF-prior LR-TF, TF-gene network information. **c,** Identification of the top 25 downstream genes (target genes) with the highest percent change in absolute expression upon blocking *NOTCH3*, *PLAUR*, *ITGA11* and *CD74*, respectively. A red highlight indicates downstream genes associated with PDAC.

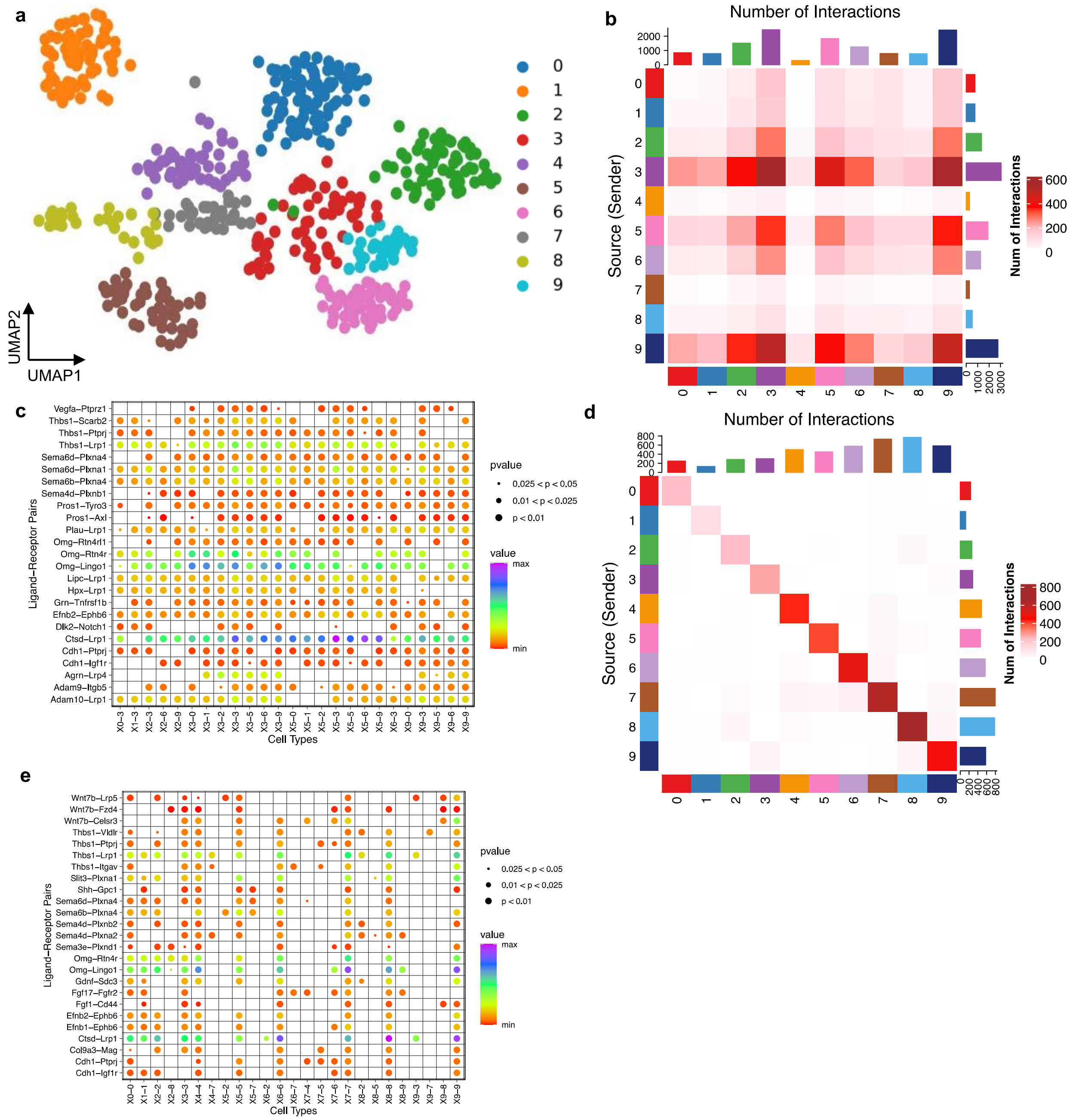


**Supplementary Figure 10: Inferred interactions by CellAgentChat on Mouse Somatosensory cortex a,** UMAP visualization of scRNA-seq data of SeqFISH+ mouse somatosensory cortex dataset. (**b, d,**) Heatmaps display the communication network inferred by CellAgentChat with non-spatial and spatial data, respectively. (**d, f,**) Dot plots illustrate the top 25 LR pairs between cell types inferred by CellAgentChat with non-spatial and spatial data, respectively.

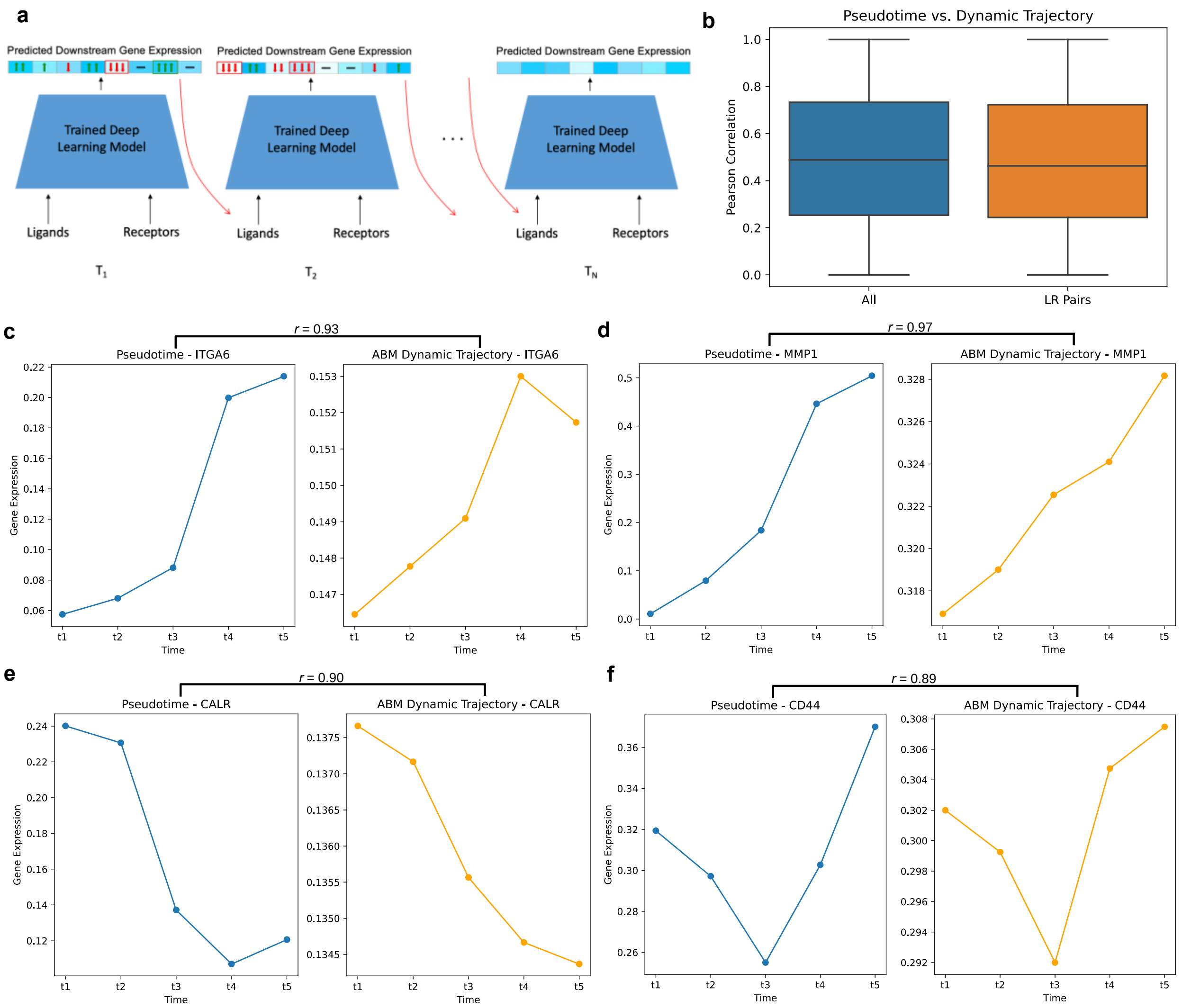


**Supplementary Figure 11: CellAgentChat Dynamic Simulations. a,** Overview of ABM dynamic update over time steps using the deep learning model. **b**, Boxplot illustrating the Pearson correlation between the pseudotime trajectory and the dynamic trajectory generated by the ABM for all genes (median $r^2$=0.49), as well as for ligand and receptor genes specifically (median $r^2$=0.48). **c,** Pseudotime trajectory vs. ABM dynamic trajectory of the receptor *ITGA6*. **d**, Pseudotime trajectory vs. ABM dynamic trajectory of the ligand *MMP1*. **e**, Pseudotime trajectory vs. ABM dynamic trajectory of the ligand *CALR*. **f**. Pseudotime trajectory vs. ABM dynamic trajectory of the receptor *CD44*.

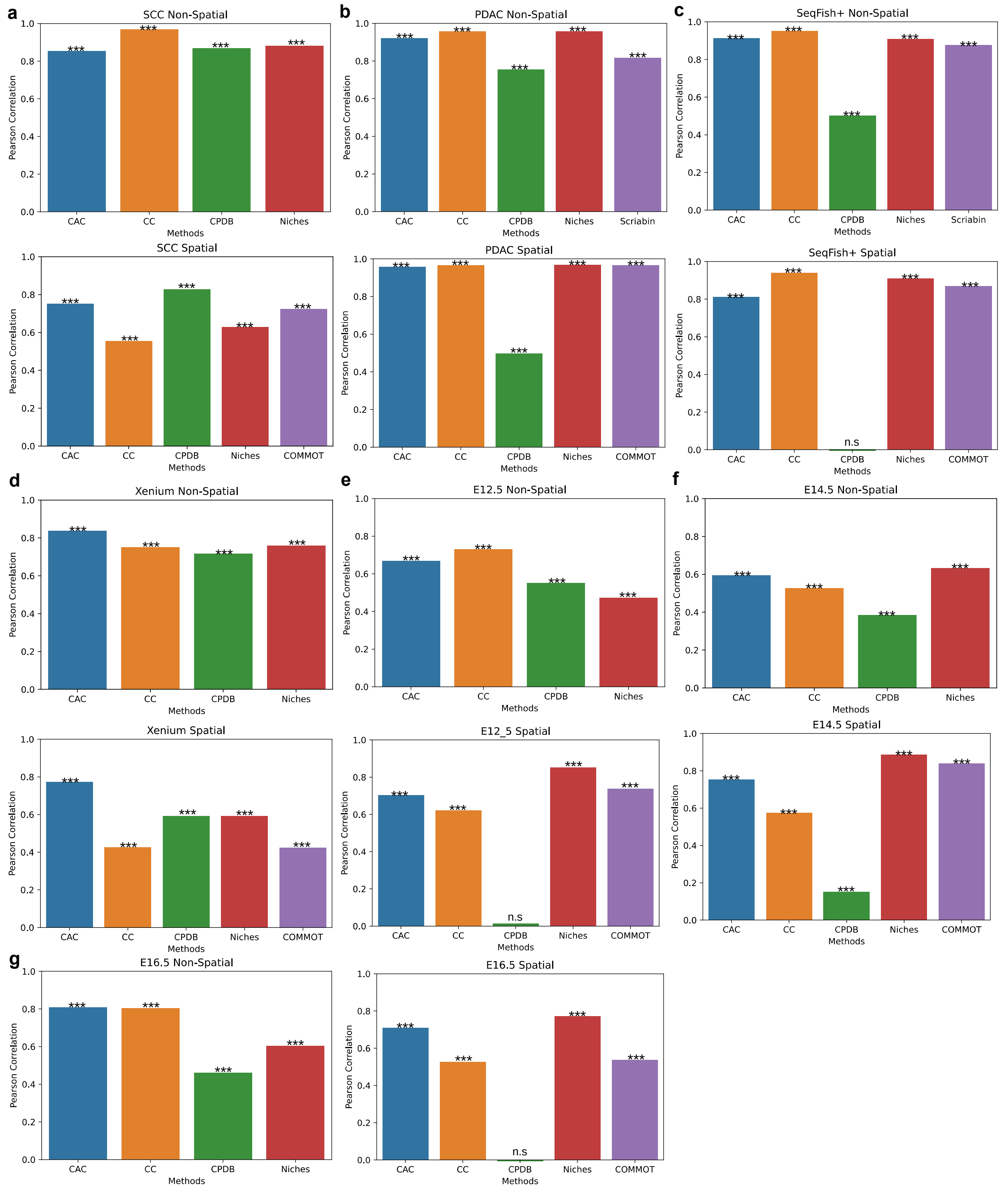


**Supplementary Figure 12: Communication pattern benchmarking of CellAgentChat with other state-of-the-art methods.** Pearson Correlation of each methods communication network compared to the ensemble communication network of all methods for the: **a,** SCC dataset. **b,** PDAC dataset. **c,** SeqFISH+ mouse cortex dataset. **d,** Xenium breast cancer dataset. **e,** E12.5 developing mouse Stereo-seq dataset. **f,** E14.5 developing mouse Stereo-seq dataset. **g,** E16.5 developing mouse Stereo-seq dataset.

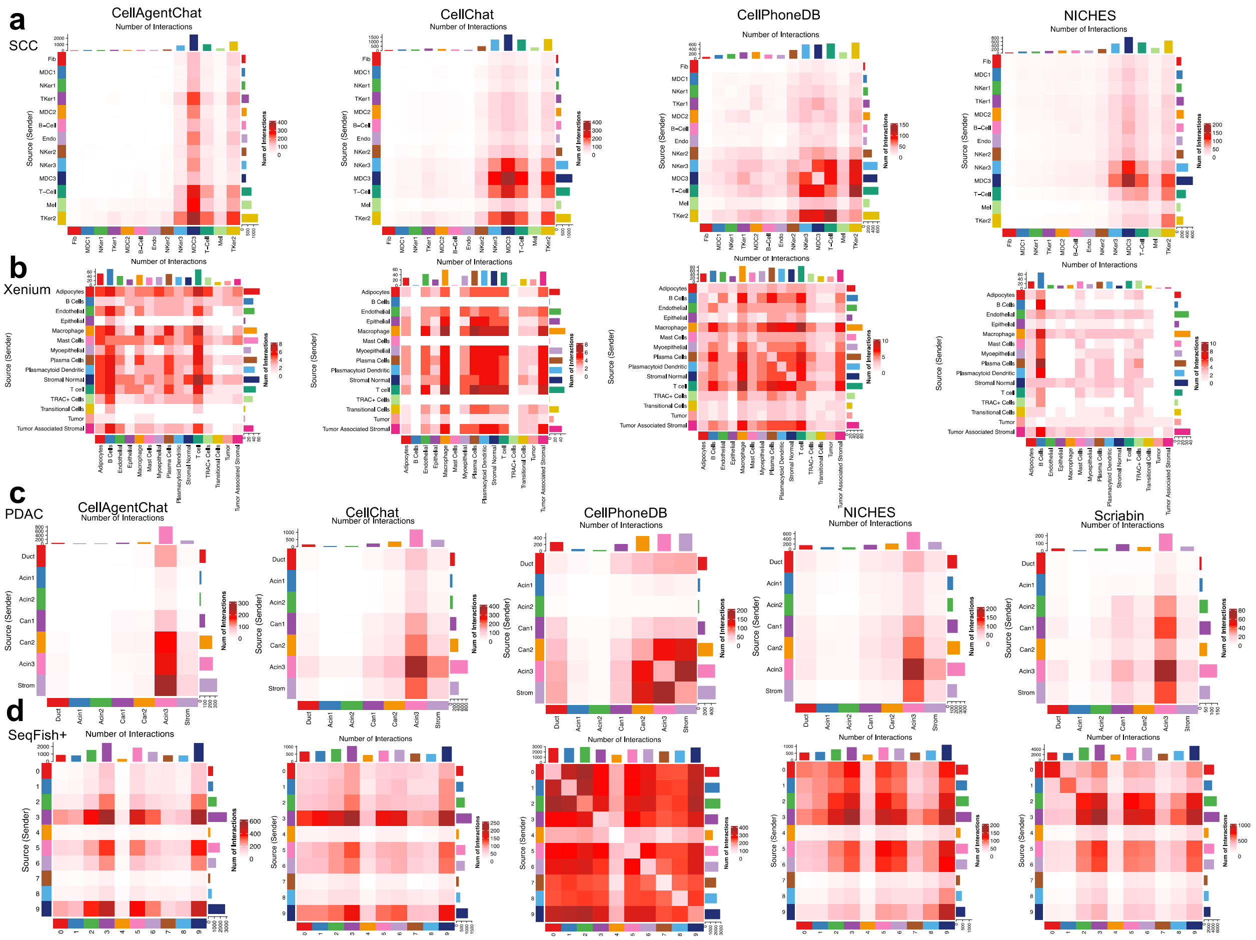


**Supplementary Figure 13: Benchmarking of CellAgentChat against other state-of-the-art methods with non-spatial data. a,** Heatmaps display the communication network between each cell type inferred by four methods for the SCC dataset. **b,** Heatmaps display the communication network between each cell type inferred by four methods for the Xenium breast cancer dataset. **c,** Heatmaps display the communication network between each cell type inferred by all five methods for the PDAC dataset. **d,** Heatmaps display the communication network between each cell type inferred by all five methods for the SeqFISH+ mouse somatosensory cortex dataset.

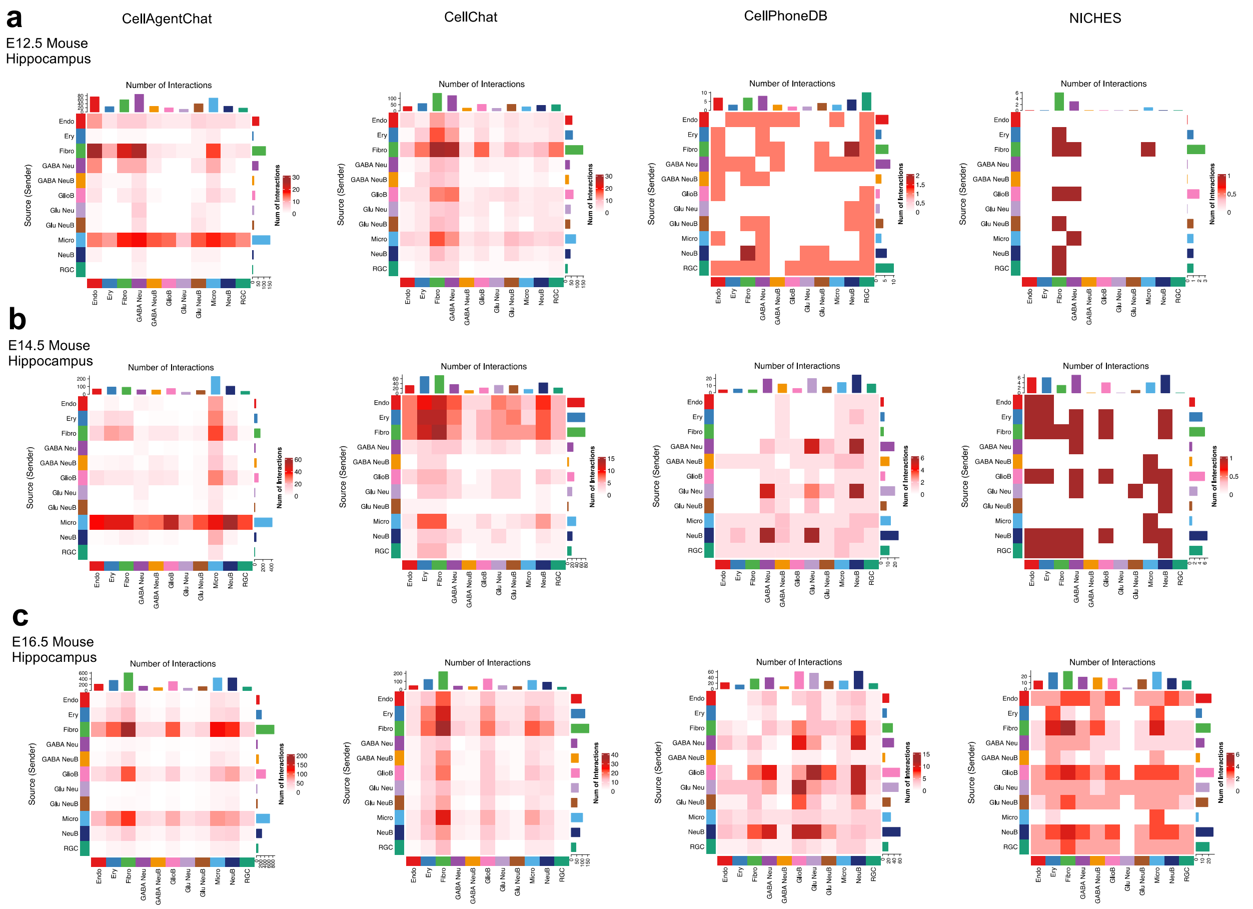


**Supplementary Figure 14: Benchmarking of CellAgentChat against other state-of-the-art methods with non-spatial data. a,** Heatmaps display the communication network between each cell type inferred by four methods for the E12.5 mouse developing hippocampus dataset. **b,** Heatmaps display the communication network between each cell type inferred by four methods for the E14.5 mouse developing hippocampus dataset. **c,** Heatmaps display the communication network between each cell type inferred by four methods for the E16.5 mouse developing hippocampus dataset.

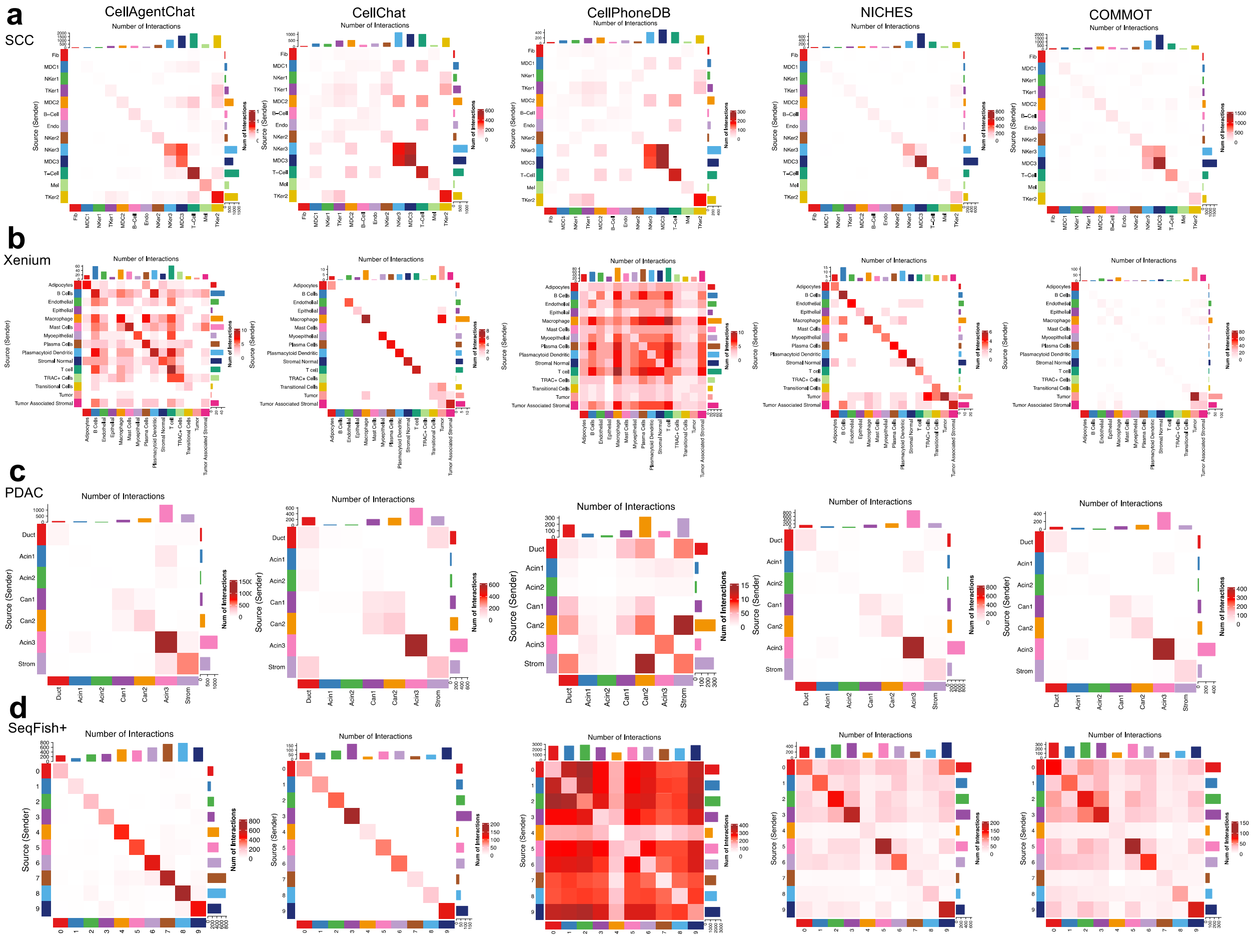


**Supplementary Figure 15: Benchmarking of CellAgentChat against other state-of-the-art methods with spatial data. a,** Heatmaps display the communication network between each cell type inferred by four methods for the SCC dataset. **b,** Heatmaps display the communication network between each cell type inferred by four methods for the Xenium breast cancer dataset. **c,** Heatmaps display the communication network between each cell type inferred by all five methods for the PDAC dataset. **d,** Heatmaps display the communication network between each cell type inferred by all five methods for the SeqFISH+ mouse somatosensory cortex dataset.

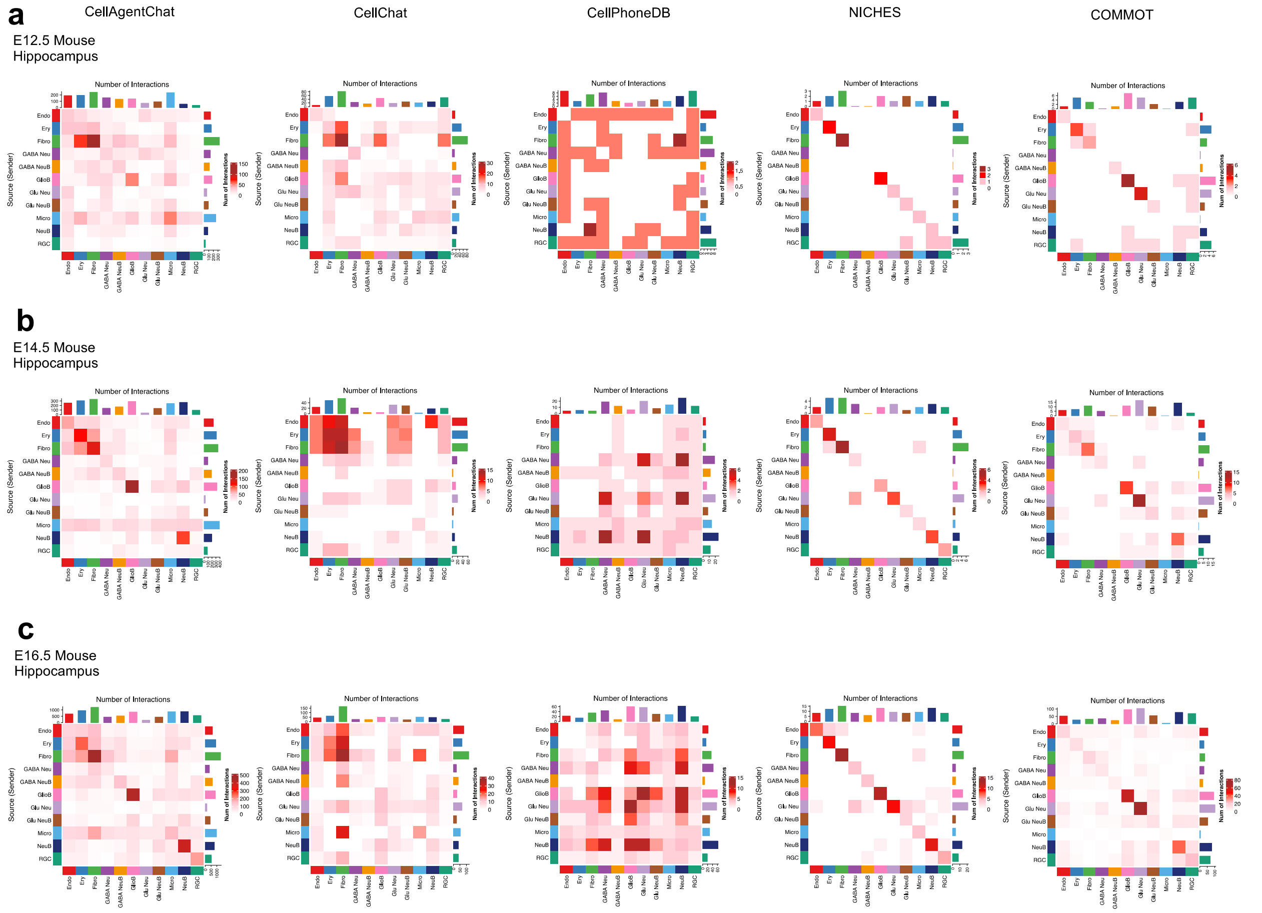


**Supplementary Figure 16: Benchmarking of CellAgentChat against other state-of-the-art methods with spatial data. a,** Heatmaps display the communication network between each cell type inferred by four methods for the E12.5 mouse developing hippocampus dataset. **b,** Heatmaps display the communication network between each cell type inferred by four methods for the E14.5 mouse developing hippocampus dataset. **c,** Heatmaps display the communication network between each cell type inferred by four methods for the E16.5 mouse developing hippocampus dataset.

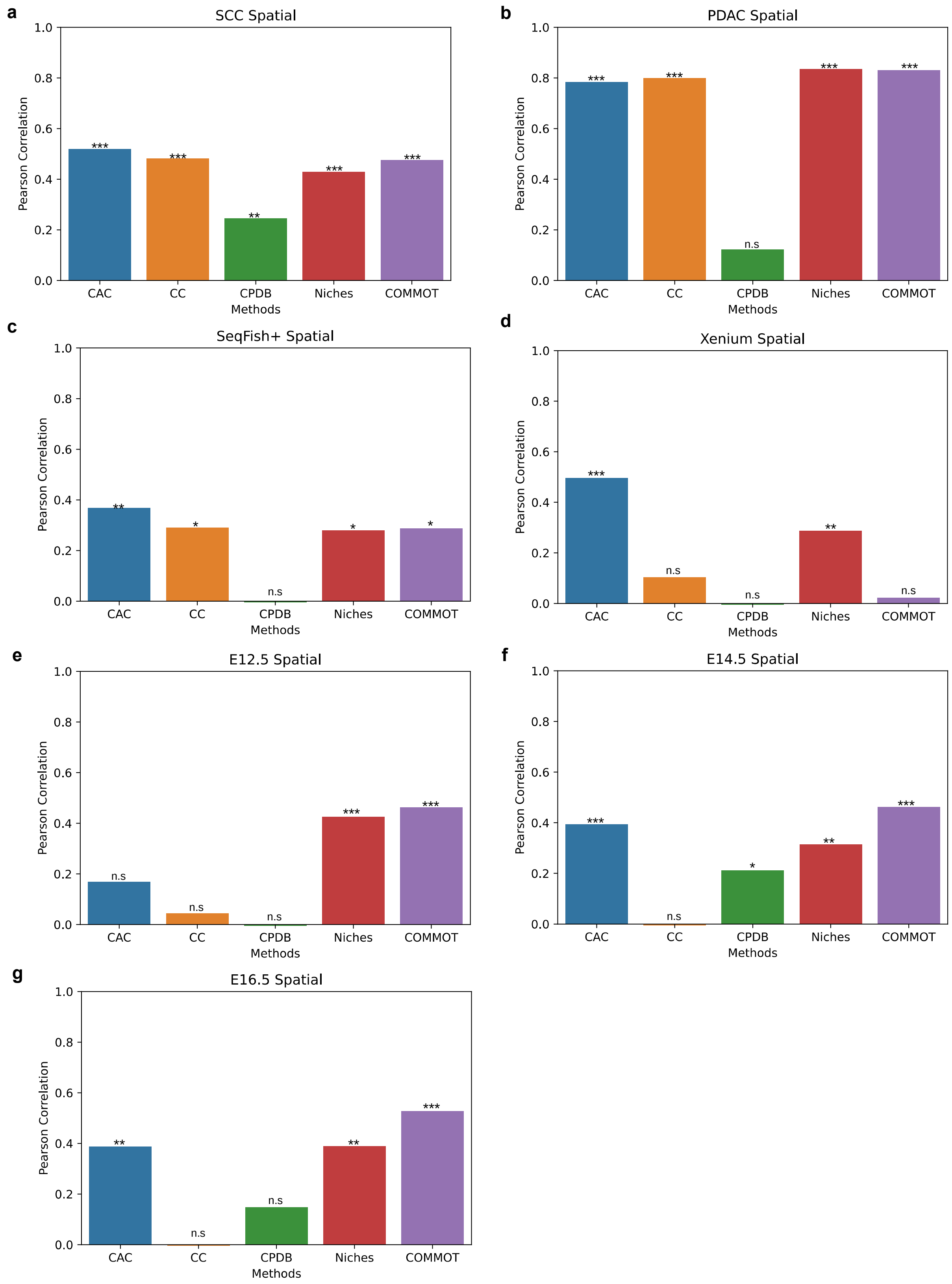


**Supplementary Figure 17: Benchmarking of CellAgentChat against other state-of-the-art methods, assessing the spatially informed nature of computed interactions.** Pearson correlation between the inferred interaction network and the inverse cell-distance matrix for the: **a,** SCC dataset. **b,** PDAC dataset. **c,** SeqFISH+ mouse cortex dataset. **d,** Xenium breast cancer dataset. **e,** E12.5 developing mouse Stereo-seq dataset. **f,** E14.5 developing mouse Stereo-seq dataset. **g,** E16.5 developing mouse Stereo-seq dataset.

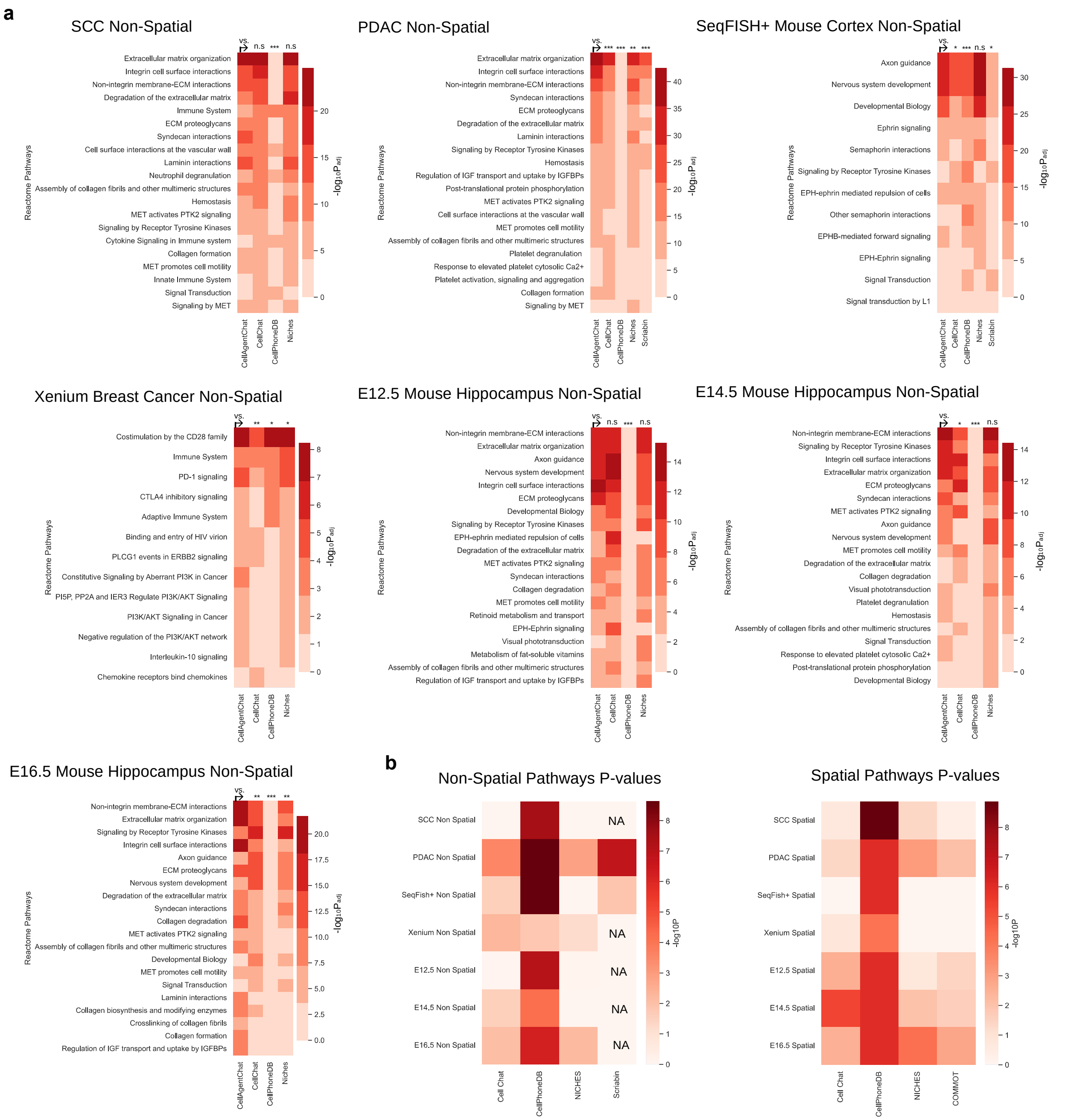


**Supplementary Figure 18: Pathway enrichment benchmarking of CellAgentChat with existing state-of-the-art methods. a,** Reactome pathway analysis conducted on receptors derived from the top 100 inferred ligand-receptor pairs identified by each method across datasets, without using spatial data. **b,** Heatmaps display the -$\log_{10}$ adjusted P-values of CellAgentChat compared to each method across datasets with non-spatial (left) and spatial (right) data.

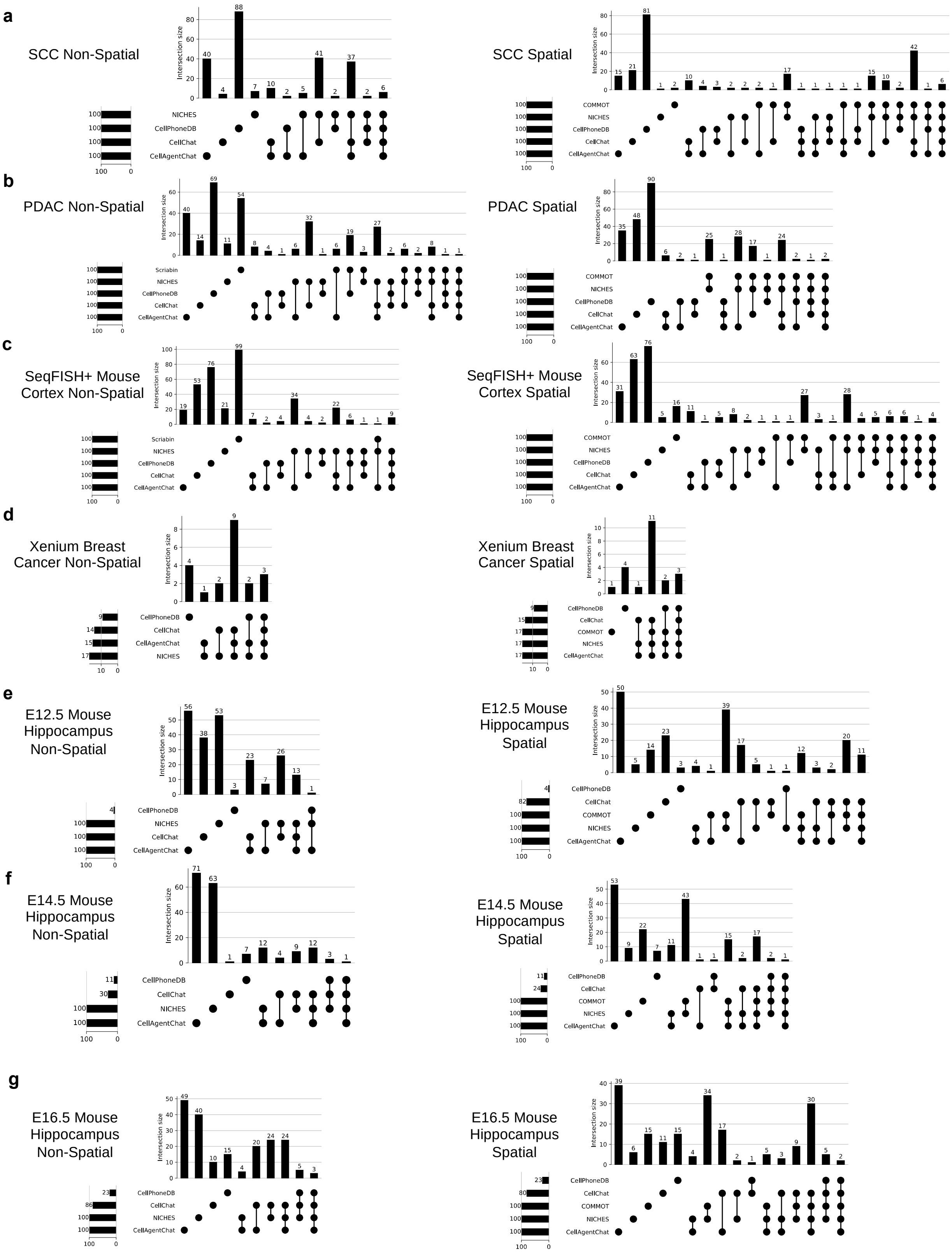


**Supplementary Figure 19: Benchmarking of CellAgentChat against other state-of-the-art methods, assessing the similarity of inferred ligand-receptor pairs.** UpSet plots show the overlap in the top 100 LR pairs inferred by each method for the: **a,** SCC dataset. **b,** PDAC dataset. **c,** SeqFISH+ mouse cortex dataset. **d,** Xenium breast cancer dataset. **e,** E12.5 developing mouse Stereo-seq dataset. **f,** E14.5 developing mouse Stereo-seq dataset. **g,** E16.5 developing mouse Stereo-seq dataset.

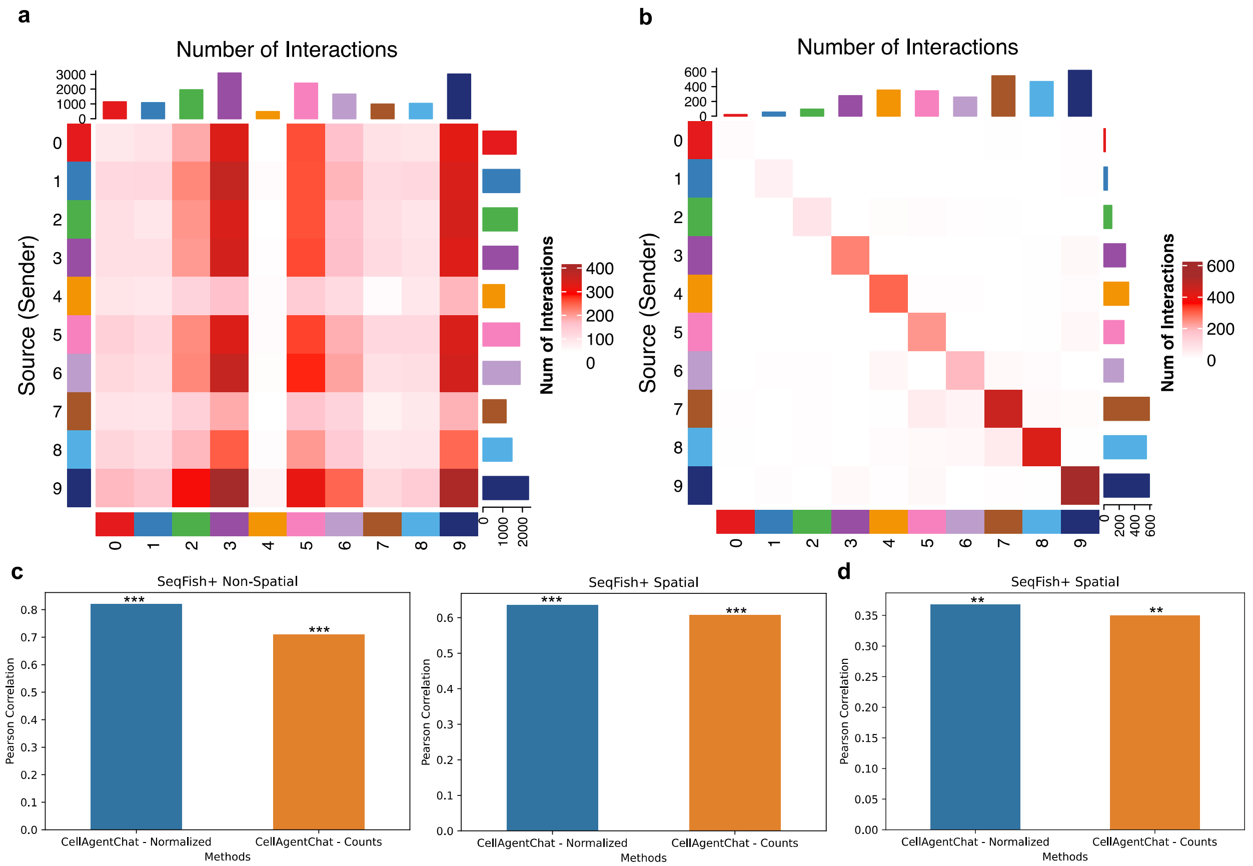


**Supplementary Figure 20: Analysis of the use of ligand and receptor raw count expression values in place of normalized and log-transformed expression values. (a, b,)** Heatmaps display the communication pattern of the SeqFISH+ mouse somatosensory cortex dataset using raw counts with non-spatial and spatial data, respectively. **c,** Pearson Correlation between inferred interaction network with normalized expression values vs raw counts and the ensemble of other state-of-the-art methods inferred interactions. **d,** Pearson Correlation between inferred interaction network with normalized expression values vs raw counts and the cell-distance matrix.

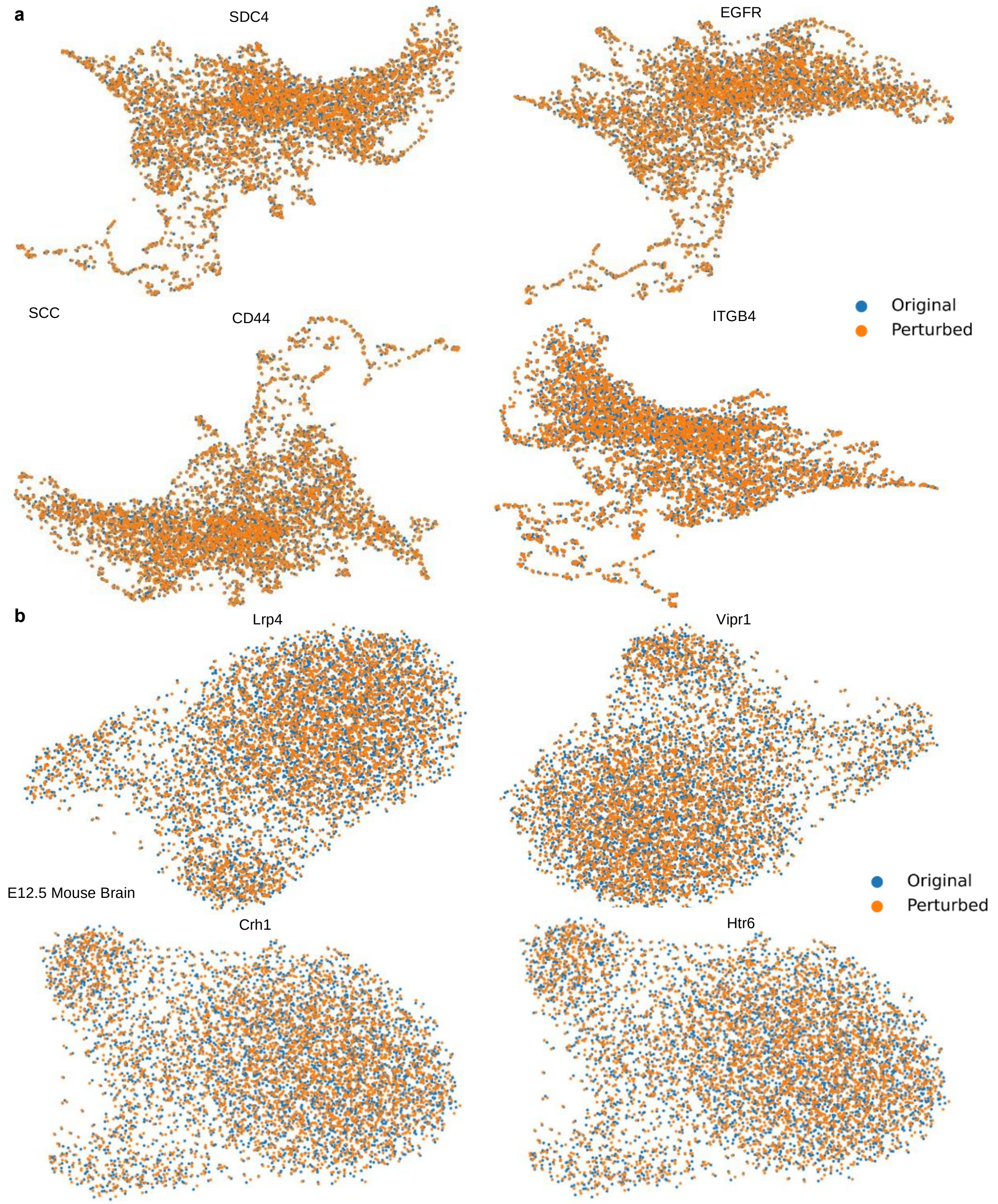


**Supplementary Figure 21: Confirmation that receptor blocking, achieved by nullifying the receptor, does not significantly alter the resulting gene expression of cells. a,** UMAP visualization of cells before and after perturbation for four receptors, *SDC4*, *EGFR*, *CD44* and *ITGB4* in the SCC dataset. UMAP projections show that the cell expression does not deviate much after receptor perturbation. **b,** UMAP visualization of cells before and after perturbation for four receptors, *Lrp1*, *Vipr1*, *Crh1* and *Htr6* in the Stereo-seq E12.5 mouse hippocampus dataset. UMAP projections show that the cell expression does not deviate much after receptor perturbation.

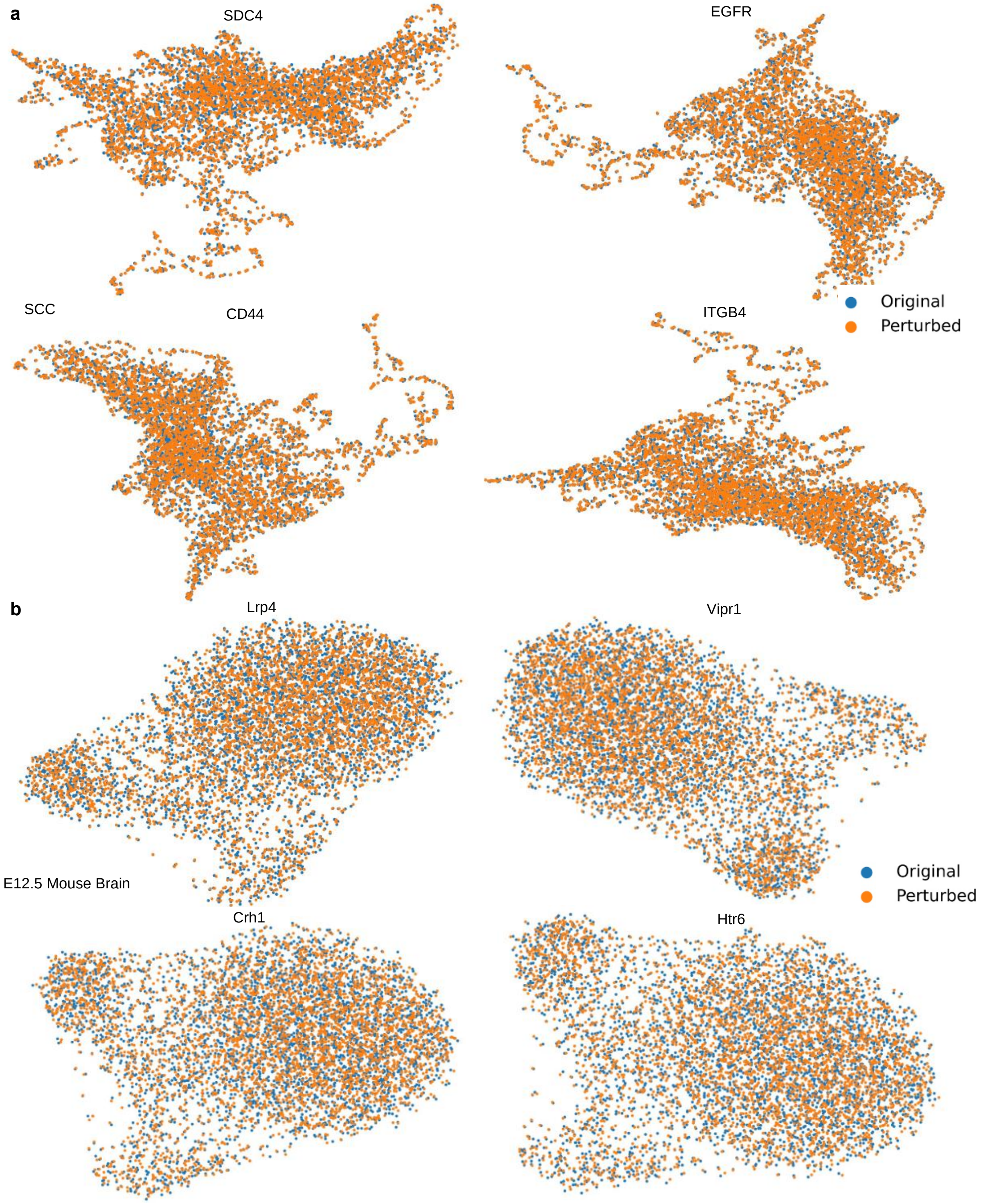


**Supplementary Figure 22: Confirmation that receptor blocking, achieved by permutating receptor features variables and lowering receptor expression, does not significantly alter the resulting gene expression of cells. a,** UMAP visualization of cells before and after perturbation for four receptors, *SDC4*, *EGFR*, *CD44* and *ITGB4* in the SCC dataset. UMAP projections show that the cell expression does not deviate much after receptor perturbation. **b,**

UMAP visualization of cells before and after perturbation for four receptors, *Lrp1*, *Vipr1*, *Crh1* and *Htr6* in the Stereo-seq E12.5 mouse hippocampus dataset. UMAP projections show that the cell expression does not deviate much after receptor perturbation.

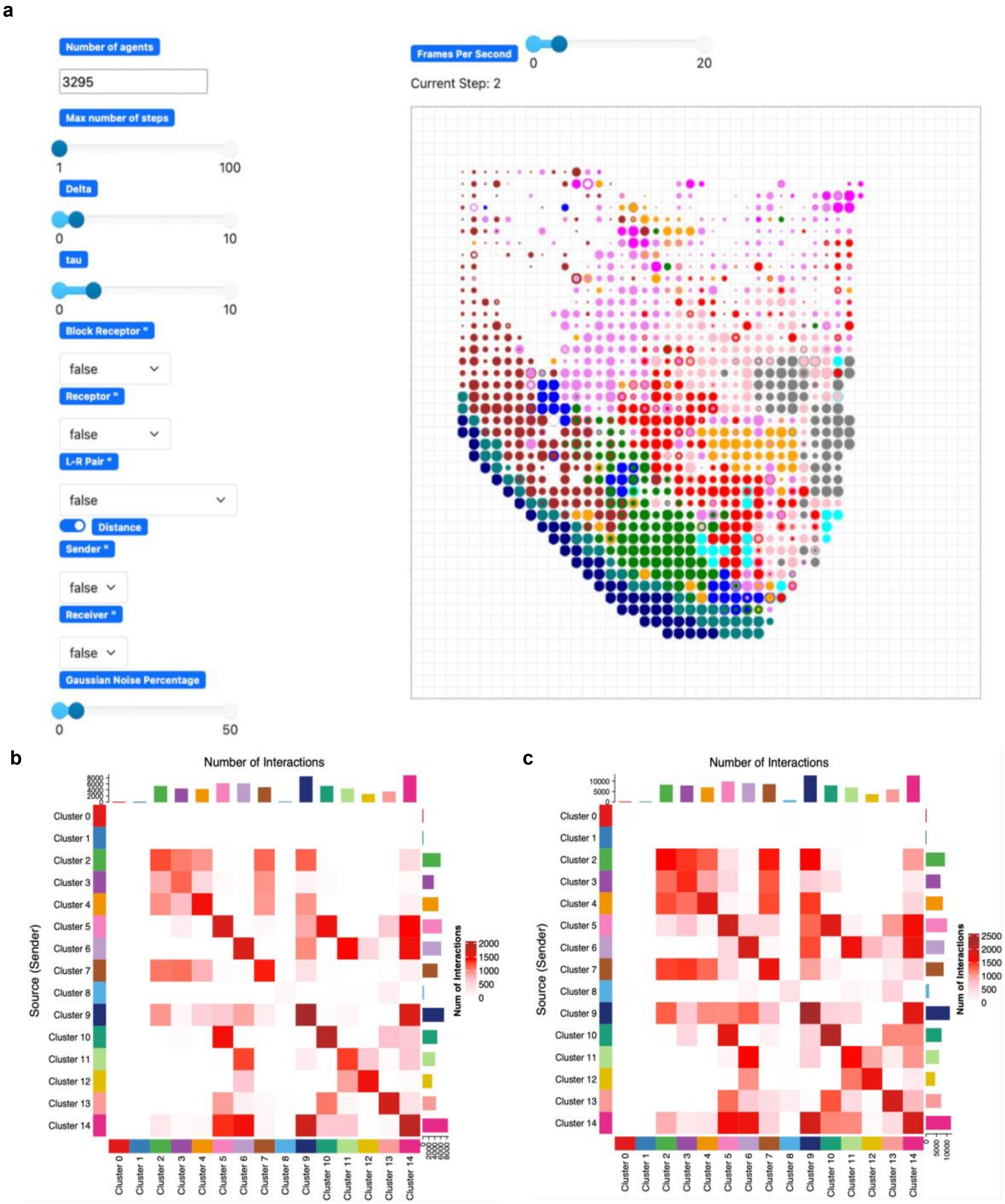


**Supplementary Figure 23: CellAgentChat animation platform and background distribution scaling for spatial data results. a,** Animation platform of CellAgentChat to visualize the changes in cell receiving score (CRS) for each cell in real-time. **b,** Heatmap of the communication network between cell types, created using significant interactions determined from the permutation test without scaling. We perform the permutation test using spatial data **c,** Heatmap of the communication network between cell types, created using significant interactions determined from the permutation test with scaling. We perform the permutation test using non-spatial data. The background gamma distribution for each ligand-receptor pair uses non-spatial mode. For scaling, we divide the scale parameter of the distribution by the average distance between all cells. Additionally, we halve both the alpha and the modified scale values.
